# Genome-wide mapping of the *Escherichia coli* PhoB regulon reveals many transcriptionally inert, intragenic binding sites

**DOI:** 10.1101/2023.02.07.527549

**Authors:** Devon Fitzgerald, Anne Stringer, Carol Smith, Pascal Lapierre, Joseph T. Wade

## Abstract

Genome-scale analyses have revealed many transcription factor binding sites within, rather than upstream of genes, raising questions as to the function of these binding sites. Here, we use complementary approaches to map the regulon of the *Escherichia coli* transcription factor PhoB, a response regulator that controls transcription of genes involved in phosphate homeostasis. Strikingly, the majority of PhoB binding sites are located within genes, but these intragenic sites are not associated with detectable transcription regulation and are not evolutionarily conserved. Many intragenic PhoB sites are located in regions bound by H-NS, likely due to shared sequence preferences of PhoB and H-NS. However, these PhoB binding sites are not associated with transcription regulation even in the absence of H-NS. We propose that for many transcription factors, including PhoB, binding sites not associated with promoter sequences are transcriptionally inert, and hence are tolerated as genomic “noise”.

**IMPORTANCE:** Recent studies have revealed large numbers of transcription factor binding sites within the genes of bacteria. The function, if any, of the vast majority of these binding sites has not been investigated. Here, we map the binding of the transcription factor PhoB across the *Escherichia coli* genome, revealing that the majority of PhoB binding sites are within genes. We show that PhoB binding sites within genes are not associated with regulation of the overlapping genes. Indeed, our data suggest that bacteria tolerate the presence of large numbers of non-regulatory, intragenic binding sites for transcription factors, and that these binding sites are not under selective pressure.

## INTRODUCTION

### Bacterial transcription factors often bind sites within genes

Bacteria encode numerous transcription factors (TFs) that regulate transcription initiation by binding DNA near promoters and modulating the ability of RNAP holoenzyme to bind promoter DNA or to isomerize to an actively transcribing conformation (1). TF function has been studied almost exclusively in the context of TF binding sites in intergenic regions, upstream of the regulated genes. However, genome-scale analyses of TF binding have identified large numbers of intragenic binding sites, far from gene starts. The proportion of binding sites for a TF that are intragenic varies extensively between different TFs (2, 3), with some TFs having the majority of their binding sites inside genes (3–6). Despite the large number of intragenic TF binding sites, relatively little is known about their function.

Regulatory activity has been described for few intragenic TF binding sites, and can be classified into distinct classes based on the regulatory target and mechanism of action: (i) canonical regulation of transcription initiation of the downstream gene, generating an RNA with an extended 5’ UTR that overlaps a gene (5, 7–10); (ii) canonical regulation of transcription initiation of a stable non-coding RNA that initiates inside a gene or 3’ UTR (11, 12); (iii) regulation of transcription initiation of the gene that contains the TF binding site; mechanisms of regulation in almost all such cases are unknown (3), although transcription repression can occur from a site close to the promoter due to a physical interaction with a more upstream site, resulting in formation of a DNA loop (13, 14); (iv) regulation of transcription elongation due to the TF acting as a road-block for RNAP (15–18). Another possible regulatory function for intragenic TF binding sites is regulation of pervasive transcription – transcription of large numbers of short, unstable RNAs from inside genes that is ubiquitous in bacteria (19, 20). Although there are no described examples of TFs that regulate unstable, intragenic transcripts, many of these RNAs are differentially expressed between growth conditions (21), consistent with regulation by TFs. Intragenic TF binding sites might also have functions that are not directly connected to gene regulation, such as facilitating short- or long-range chromosome contacts (22–25), or serving as TF-titrating decoy sites (26, 27). Lastly, it is possible that intragenic TF binding sites serve no biological function, and arise as a consequence of genetic drift (28), or genome evolution that is constrained by selection for particular codons.

### PhoB is a conserved transcription factor that regulates phosphate homeostasis

PhoB is a member of the PhoB/OmpR family of response regulator TFs, and is a key regulator of phosphate homeostasis in many Gram-negative bacteria (29, 30). PhoB forms a two-component system with the sensor kinase PhoR (31). When inorganic phosphate (P_i_) levels are low, PhoR autophosphorylates and then phosphorylates PhoB (30, 31), triggering PhoB dimerization and DNA binding activity (30). Phosphorylated PhoB binds direct repeat sequences called *pho* boxes (32), and is a dual regulator, capable of both activating and repressing transcription depending on the position of the binding site.

In *Escherichia coli* and related species, PhoB regulates expression of genes encoding the high affinity phosphate transport system (*pst*), a phosphonate transport complex (*phn*), the glycerol-3-phosphate transporter (*ugp*), and other genes related to phosphate homeostasis (30, 33). These genes are collectively referred to as the *pho* regulon. PhoB has been implicated in regulation of a number of other cellular processes and stress responses, including motility, biofilm formation, quorum sensing, cell surface remodeling, stringent response, and the general stress response (34, 35). Indeed, transcriptomic and proteomic studies of phosphate-depleted *E. coli* have suggested that the *pho* regulon has many additional members (36, 37). However, most of these putative regulon members have limited experimental support (30, 33).

Here, we describe a high-resolution, genome-wide mapping of the *pho* regulon using ChIP-seq and RNA-seq. We refine and expand the set of known *pho* regulon genes and identify many intragenic PhoB binding sites. We show that the large majority of intragenic PhoB binding sites are not conserved, and are not associated with detectable regulatory function. Thus, our data suggest that individual intragenic PhoB sites are non-functional, and that TFs can bind many intragenic sites with little or no impact on local transcription.

## RESULTS

### Genome-wide Binding of PhoB under phosphate-limiting conditions

ChIP-seq is used to map the genome-wide binding of TFs. To facilitate ChIP-seq of *E. coli* PhoB, we introduced C-terminal FLAG tags at the native *phoB* locus. We used quantitative reverse-transcriptase PCR (qRT-PCR) to measure expression of *pstS*, a PhoB-activated gene, in wild-type cells, Δ*phoB* cells, and cells expressing *phoB*-FLAG_3_. Cells were grown in minimal medium with low phosphate levels, to induce the kinase activity of PhoR. As expected, we observed a large decrease (∼900-fold) in *pstS* levels in Δ*phoB* cells relative to wild-type cells (Figure 1). In cells expressing PhoB-FLAG_3_, we observed a much smaller decrease (∼8-fold) in *pstS* levels relative to wild-type cells (Figure 1), indicating that the tagged PhoB derivative retains partial function.

**Figure 1.**
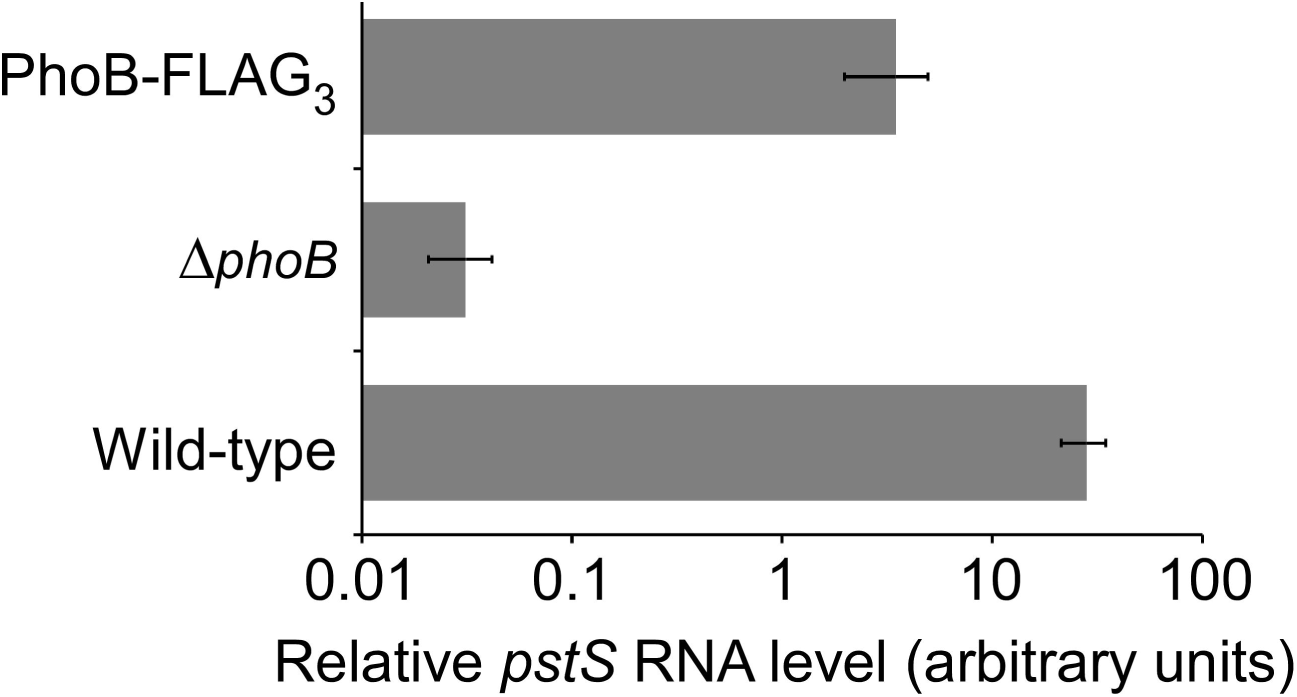
Partially reduced activity of the C-terminally FLAG_3_-tagged PhoB. Quantitative RT-PCR was used to measure levels of the *pstS* RNA relative to the *minD* RNA control in wild-type MG1655 + pBAD24 (“Wild-type”), MG1655 Δ*phoB* (CDS091) + pBAD24 (“Δ*phoB*”), or MG1655 *phoB*-FLAG_3_ (DMF34) + pBAD24 (“PhoB-FLAG_3_”), for cells grown under low phosphate conditions. Values represent the average of three independent biological replicates; error bars represent +/- one standard deviation.

We used ChIP-seq to map the genome-wide binding of PhoB-FLAG_3_ during growth under low phosphate conditions. Thus, we identified 65 enriched regions (Figure 2A; Table 1). As a control, we performed ChIP-seq with an untagged strain grown under the same conditions; none of the regions enriched in the PhoB-FLAG_3_ ChIP-seq dataset were enriched in the control dataset (Figure 2A). We conclude that the 65 enriched regions in the PhoB-FLAG_3_ ChIP-seq dataset are likely to represent genuine PhoB-bound regions. Note that a single PhoB-bound region could include more than one PhoB site, as is the case for the region upstream of *phoB* itself, which has been reported to include two PhoB sites (38). We identified a highly enriched sequence motif, with instances of the motif found in 59 of the 65 putative PhoB-bound regions (MEME *E*-value = 3.0e^-72^; Figure 2B). This motif contains a clearly distinguishable direct repeat and is similar to the previously reported *pho* box consensus sequence (39). Furthermore, the identified motif is centrally enriched relative to the calculated ChIP-seq peak centers (Figure 2C; Centrimo *E*-value = 1.2e^-12^). The presence and central enrichment of this motif at ChIP-seq peaks further supports the veracity of PhoB-bound regions and confirms the high spatial resolution of the ChIP-seq data.

**Figure 2.**
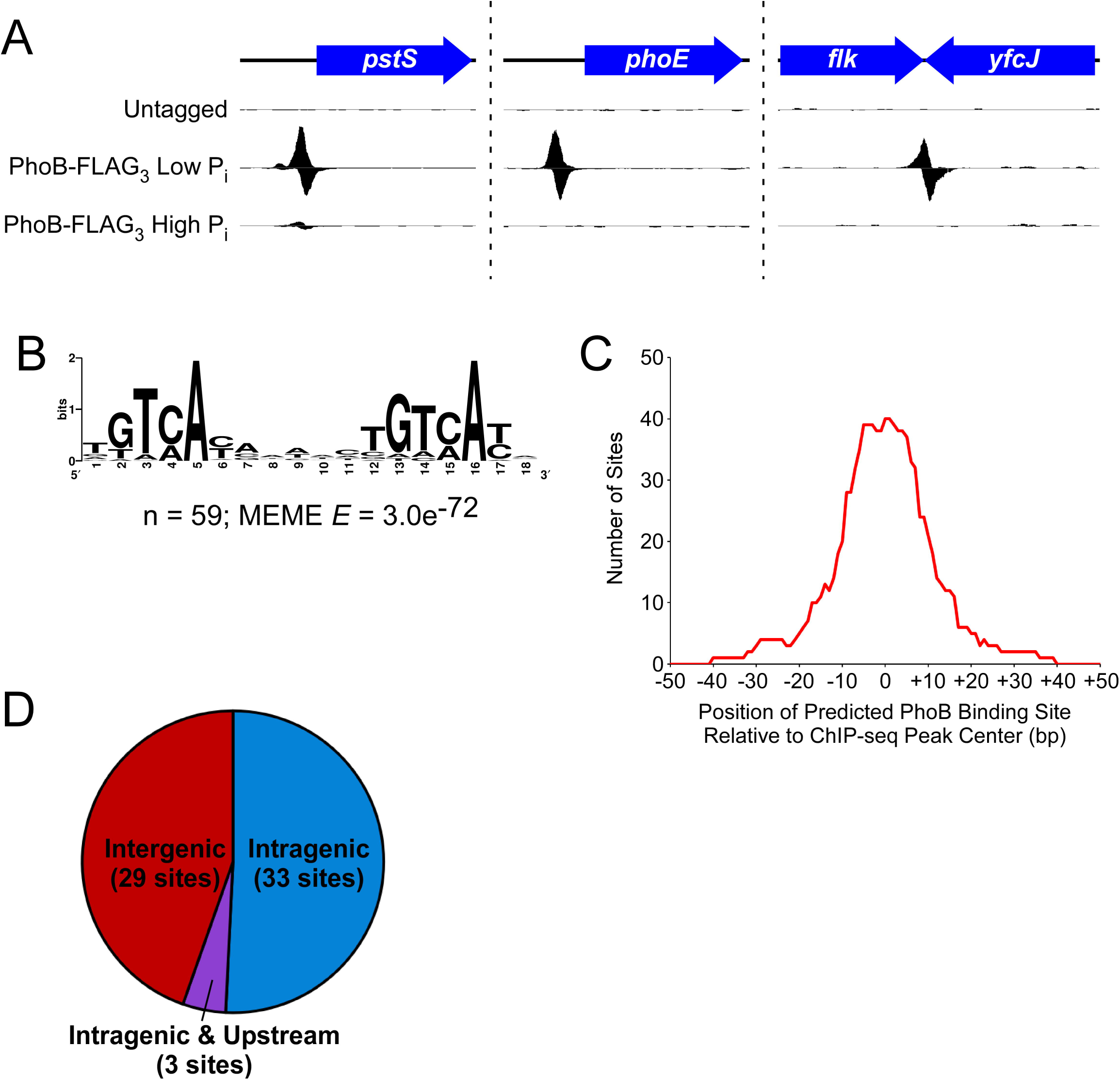
ChIP-seq identifies PhoB binding sites. **(A)** ChIP-seq data for (i) an untagged control under low phosphate conditions, (ii) PhoB-FLAG_3_ under low phosphate conditions, and (iii) PhoB-FLAG_3_ under high phosphate conditions. Three genomic regions are shown, with one dataset from two independent biological replicates. Values on the *x*-axis represent genome position. Values on the *y*-axis represent normalized sequence read coverage, with positive values indicating sequence reads mapping to the forward strand, and negative values indicating sequence reads mapping to the reverse strand. *y*-axis scales differ between the three genomic regions but are matched for the three datasets for any given genomic region. **(B)** Significantly enriched DNA sequence motif derived from 100 bp regions surrounding each ChIP-seq peak. The number of sites contributing to the motif and the *E*-value determined by MEME are indicated. **(C)** Analysis of the position of inferred PhoB binding sites relative to the position of ChIP-seq peak centers. For each of the binding sites contributing to the motif determined by MEME (see panel (C)), we determined the position of the binding site relative to the associated ChIP-seq peak center. The *x*-axis indicates position relative to ChIP-seq peak centers. The *y*-axis indicates the number of binding sites that cover any given position. **(D)** Pie-chart showing the genome context of PhoB binding sites identified by ChIP-seq. Sites designated as “Intragenic & Upstream” are intragenic but <200 bp upstream of an annotated gene start.

**Table 1.**
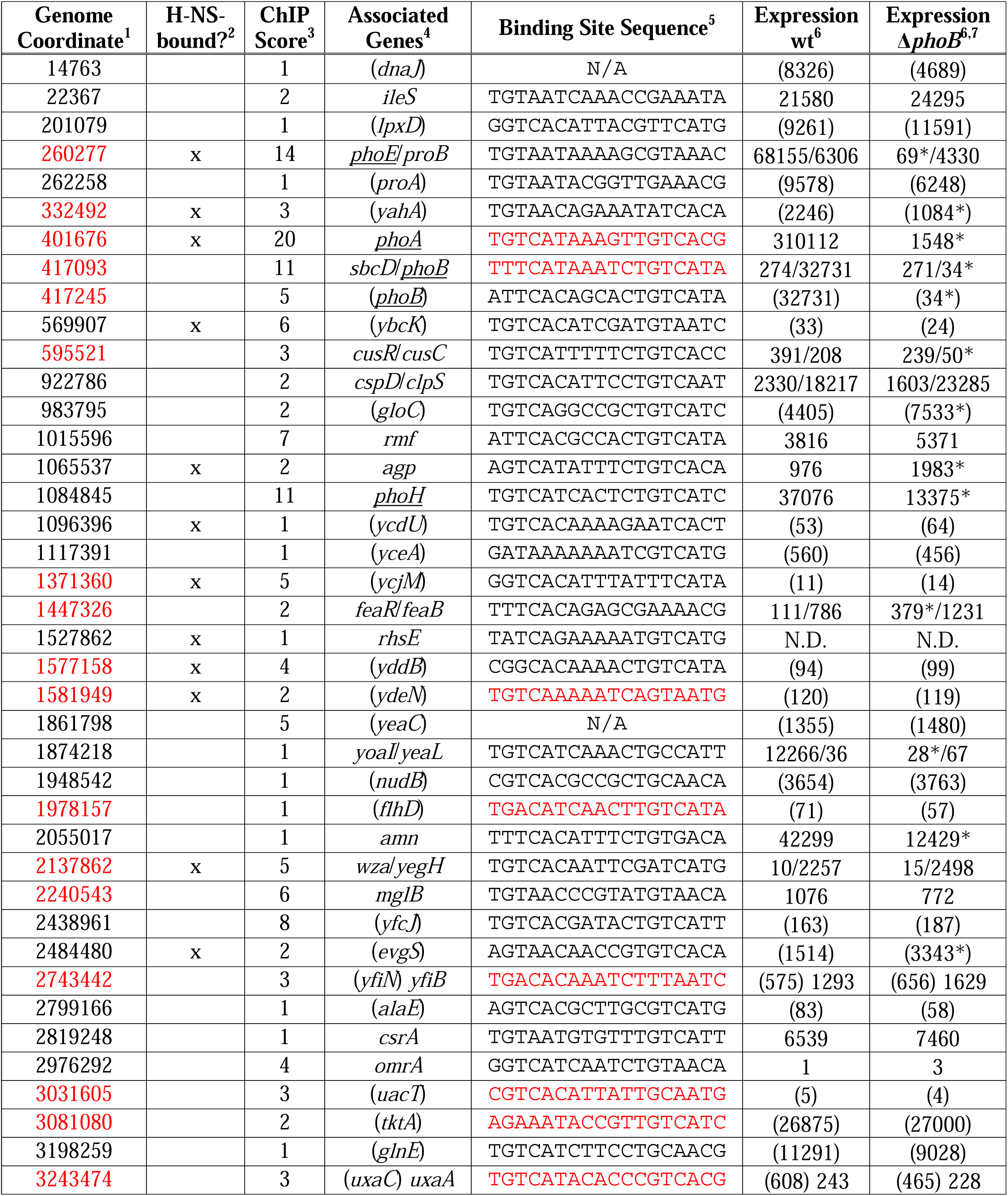
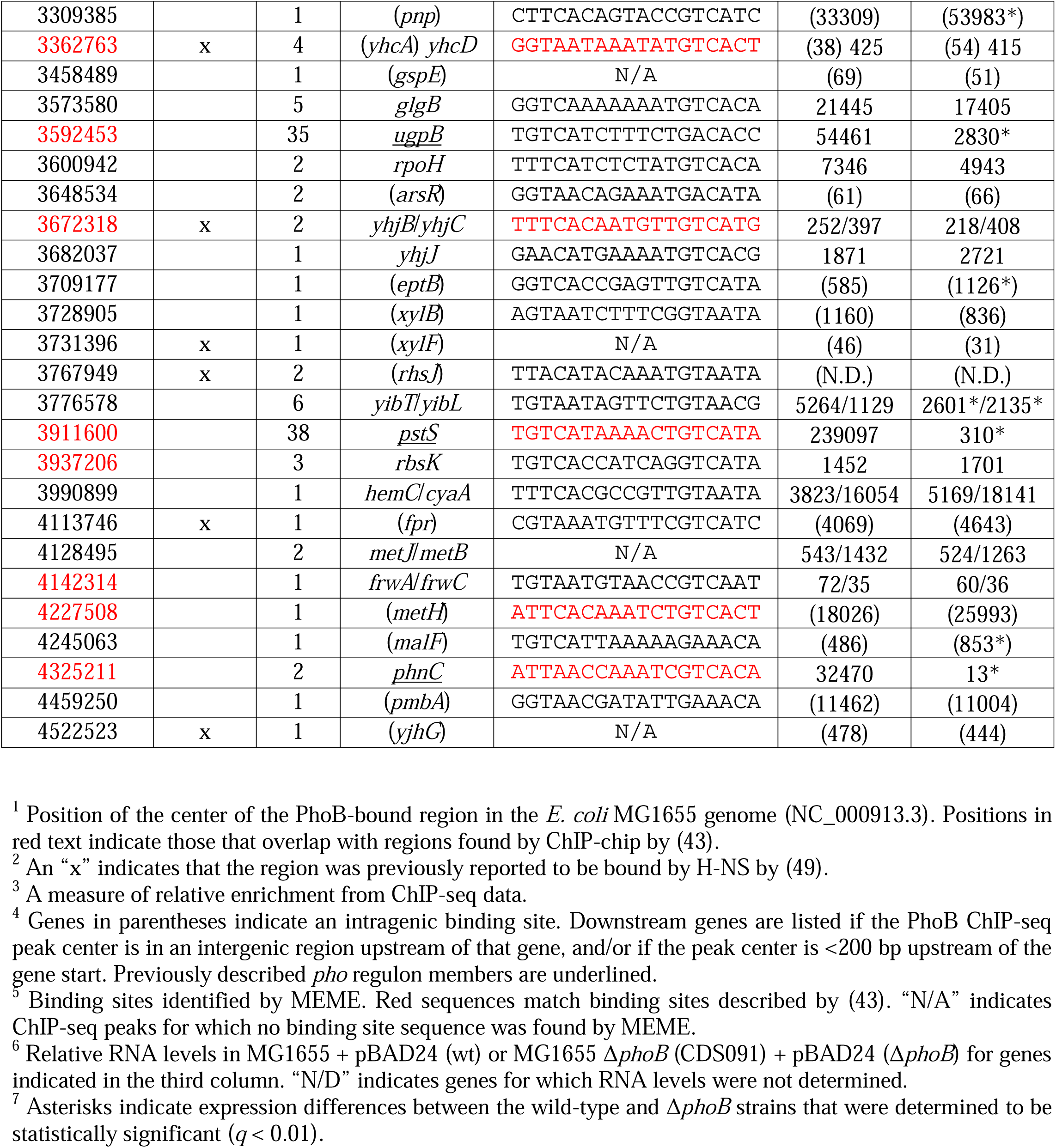
List of PhoB-bound regions identified by ChIP-seq.

The 65 PhoB-bound regions identified by ChIP-seq include most well-established PhoB sites, as well as many novel targets (Table 1). We identified PhoB-bound regions upstream of 7 of the 10 genes/operons described previously as being in the *pho* regulon (Table 1, underlined gene names) (33), with no ChIP-seq signal upstream of *waaH*, *ytfK*, or *psiE*. We also identified PhoB-bound regions upstream of the predicted *pho* regulon gene *amn* (Table 1) (40), and upstream of *yoaI*, which was described as a direct PhoB target in *E. coli* O157:H7 (41, 42). We identified 21 PhoB-bound regions upstream of genes/operons not previously described as part of the *pho* regulon, and lacking a clear connection to phosphate homeostasis. The remaining 36 PhoB-bound regions, over half of the total sites identified by ChIP-seq, are located inside genes (Figure 2D; Table 1). Strikingly, all but 3 intragenic PhoB binding sites are far from neighboring gene starts (>200 bp), thus are unlikely to participate in promoter-proximal regulation of these genes (Figure 2D; Table 1).

Our PhoB ChIP-seq data show only modest agreement with an earlier study that identified many putative PhoB binding sites using ChIP-chip (43), although both studies are consistent in the lack of signal upstream of *waaH*, *ytfK*, or *psiE*. Of the 43 ChIP-chip peaks identified by Yang *et al*., 24 are <400 bp from a ChIP-seq peak in our data (Table 1, coordinates in red), while the remaining 19 ChIP-chip peaks are >2,800 bp from the closest ChIP- seq peak. Even for the 24 ChIP-chip peaks close to ChIP-seq peaks identified in the current study, the peak centers calculated from the two datasets are up to 383 bp apart, and only 13 regions share a motif call between studies (Table 1, motifs in red). These discrepancies between datasets are probably due, at least in part, to the low resolution of the ChIP-chip data, and differences in peak-calling and motif-calling algorithms. It is challenging to determine whether peak calls from each of the two studies represent the same biological binding events. We performed *de novo* motif identification for three sets of peaks: (i) shared, (ii) unique to the current study, and (iii) unique to Yang *et al*. For (i) shared peaks and (ii) peaks unique to the current study, 100 bp of sequence surrounding each ChIP-seq peak center was extracted and analyzed by MEME. In both cases, highly enriched sequence motifs were found that are close matches to the expected PhoB motif (Figure 3A + B). For (iii) sites unique to the Yang *et al.* dataset, the same analysis was performed using both 100 bp and 500 bp windows surrounding the published peak center locations. The resulting motifs were poorly enriched (MEME *E*-values >1) and bear no similarity to the expected PhoB motif. We conclude that most or all of the 40 regions unique to the current study represent genuine PhoB binding sites, while those unique to the Yang *et al.* study largely do not.

**Figure 3.**
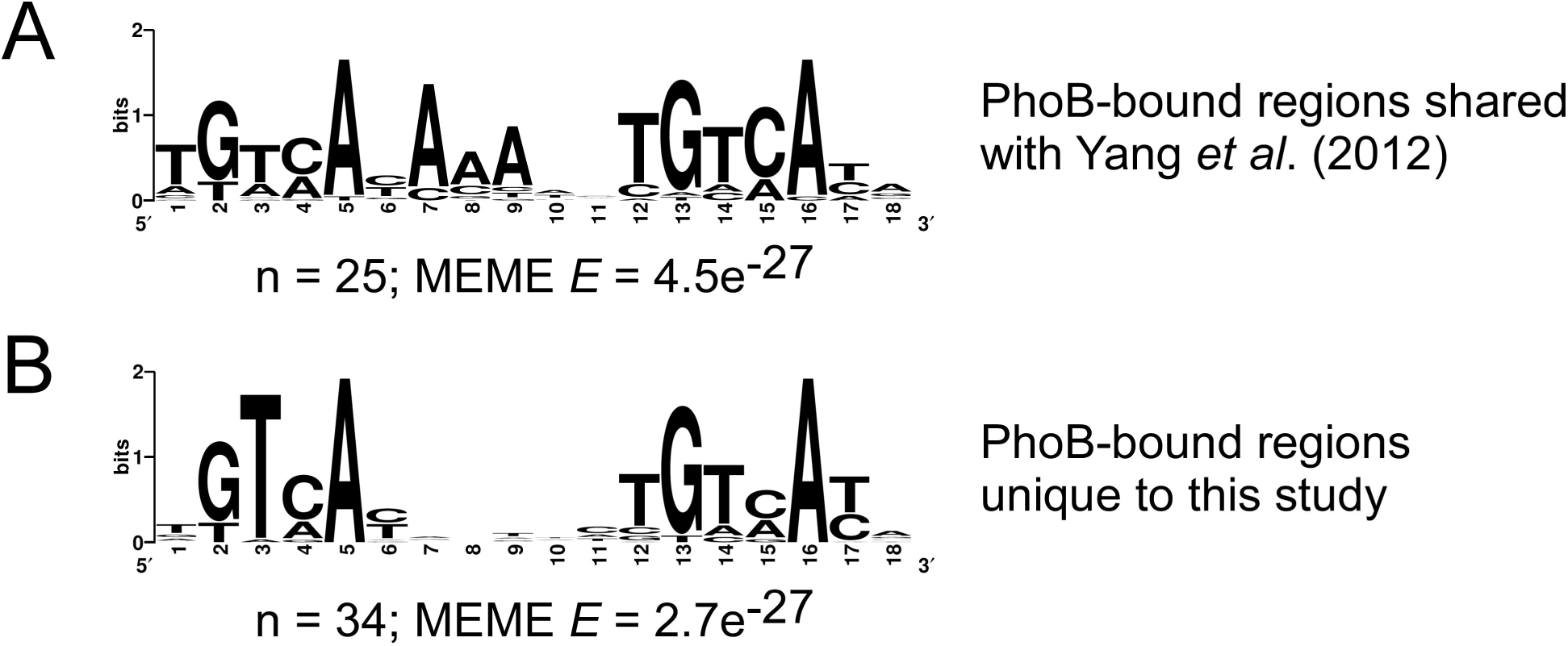
Comparison of ChIP-seq and ChIP-chip datasets. **(A)** Significantly enriched DNA sequence motif derived from 100 bp regions surrounding each ChIP-seq peak for regions shared between the ChIP-seq dataset and a published ChIP-chip dataset (43). The number of sites contributing to the motif, and the *E*-value determined by MEME are indicated. **(A)** Significantly enriched DNA sequence motif derived from 100 bp regions surrounding each ChIP-seq peak for regions unique to the ChIP-seq dataset, i.e. not found in the published ChIP-chip dataset (43). The number of sites contributing to the motif, and the *E*-value determined by MEME are indicated.

### Genome-wide Binding of PhoB under high phosphate conditions

To determine whether PhoB binds any target DNA sites when PhoR is inactive, we repeated the ChIP-seq experiment, but grew cells under conditions with high phosphate. We detected only a single PhoB-bound region: the intergenic region upstream of *pstS* (Figure 2A). This is a well-established site of PhoB binding, and was the most enriched PhoB-bound region in the low phosphate ChIP-seq experiment. As expected, PhoB binding upstream of *pstS* was substantially lower under conditions of high phosphate than under conditions of low phosphate (Figure 2A). Thus, our data suggest that under conditions of high phosphate, PhoB weakly regulates *pstS*, but does not regulate any of its other target genes.

### Reassessing the pho regulon

To address whether the detected PhoB sites contribute to transcription regulation, RNA-seq was performed using wild-type and Δ*phoB* strains grown in low-phosphate medium. In total, 181 genes were differentially expressed between the wild-type and the Δ*phoB* strains (*p*-value ≤ 0.01, >2-fold difference in RNA levels; Figure 4; Table S1). We observed significant positive regulation of all 7 reported *pho* regulon operons for which we observed upstream PhoB binding by ChIP-seq, i.e., *phnCDEFGHIJKLMNOP*, *phoH*, *ugpBAECQ*, *pstSCAB*- *phoU*, *phoA*-*psiF*, *phoE*, and *phoBR* (Table 1 + S1; positive regulation was observed for all genes in all operons, except for *phoB* which could not be assessed in the Δ*phoB* strain). We also observed significant positive regulation of *amn* and *yoaI*; ChIP-seq identified PhoB binding upstream of these genes, and although they have not generally been considered as part of the *pho* regulon, they have been previously reported as being direct PhoB targets (41, 42).

**Figure 4.**
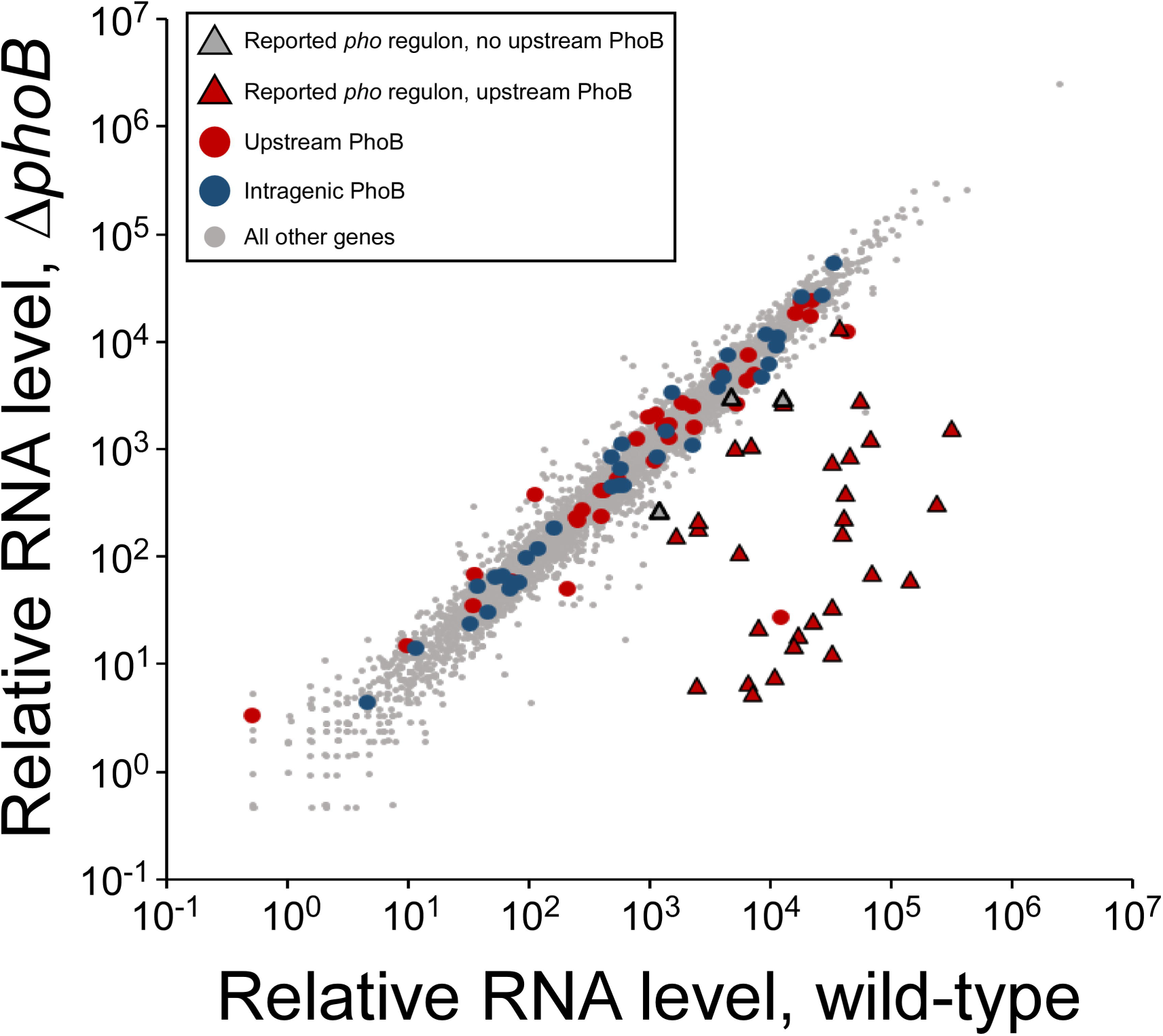
RNA-seq analysis of wild-type and. Δ***phoB E. coli*.** Scatter-plot showing relative RNA levels for all genes in wild-type (MG1655 + pBAD24) or Δ*phoB* (CDS091 + pBAD24) cells. Each datapoint corresponds to a gene. Triangle datapoints represent genes previously reported to be in the *pho* regulon, with red fill indicating that the transcript has an upstream PhoB site identified by ChIP-seq, and gray fill indicating no upstream site. Red circle datapoints represent genes not previously reported to be in the *pho* regulon but with upstream PhoB sites identified by ChIP-seq. Blue circle datapoints represent genes with internal PhoB sites identified by ChIP- seq. All other genes are represented by gray circle datapoints.

We observed significant positive regulation of *ytfK* and *waaH*, reported *pho* regulon genes that lack associated PhoB binding. We conclude that *ytfK* and *waaH* are regulated indirectly by PhoB. By contrast, we did not observe significant regulation of known and predicted *pho* regulon genes *psiE*, *asr*, *eda*, *argP*, and *pitB*; none of these genes had detectable upstream PhoB binding by ChIP-seq. We conclude that *psiE*, *asr*, *eda*, *argP*, and *pitB* are unlikely to be regulatory targets of PhoB.

To identify novel *pho* regulon genes, we determined whether any additional genes with associated PhoB binding were significantly differentially expressed between wild-type and Δ*phoB* cells. We observed significant, >2-fold positive regulation of *cusC* and *yibT*, and significant, >2-fold negative regulation of *feaR* and *agp*, genes with upstream PhoB binding as determined by ChIP-seq. We also observed significant regulation of 6 genes with internal PhoB sites: *yahA*, *gloC*, *pnp*, *evgS*, *eptB* and *malF*. Although only two of these genes (*yahA* and *evgS*) were differentially expressed >2-fold between wild-type and Δ*phoB* cells, all but *yahA* were more highly expressed in Δ*phoB* cells than wild-type cells. We hypothesized that PhoB represses transcription of these genes by acting as a roadblock for RNAP. To test this hypothesis, we grew wild-type and Δ*phoB* cells under phosphate-limiting conditions and measured RNAP (β subunit) occupancy upstream and downstream of the PhoB sites within the *gloC*, *pnp*, and *evgS* genes using ChIP-qPCR. As controls, we measured RNAP occupancy within the *pstS*, and *ugpB* genes, confirmed members of the *pho* regulon. We also measured RNAP occupancy within the *yoaI* and *amn* genes that the combined RNA-seq and ChIP-seq data suggested are members of the *pho* regulon. As expected, we observed substantially higher RNAP occupancy within *pstS* and *ugpB* in wild-type cells than in Δ*phoB* cells (Figure 5). Moreover, we observed substantially higher RNAP occupancy within *yoaI* and *amn* in wild-type cells than in Δ*phoB* cells (Figure 5), supporting the idea that these genes are part of the *pho* regulon. For the three genes with intragenic PhoB sites, we reasoned that if PhoB acts as a roadblock to elongating RNAP, RNAP occupancy downstream of PhoB sites would increase in Δ*phoB* cells relative to that in wild-type cells and relative to any change in RNAP occupancy upstream of the PhoB site. However, we did not observe significant increases in relative RNAP occupancy downstream of PhoB sites for any of the three genes (Figure 5; *p* > 0.2; see Methods for details of the statistical analysis), suggesting that PhoB regulation of these genes is indirect.

**Figure 5.**
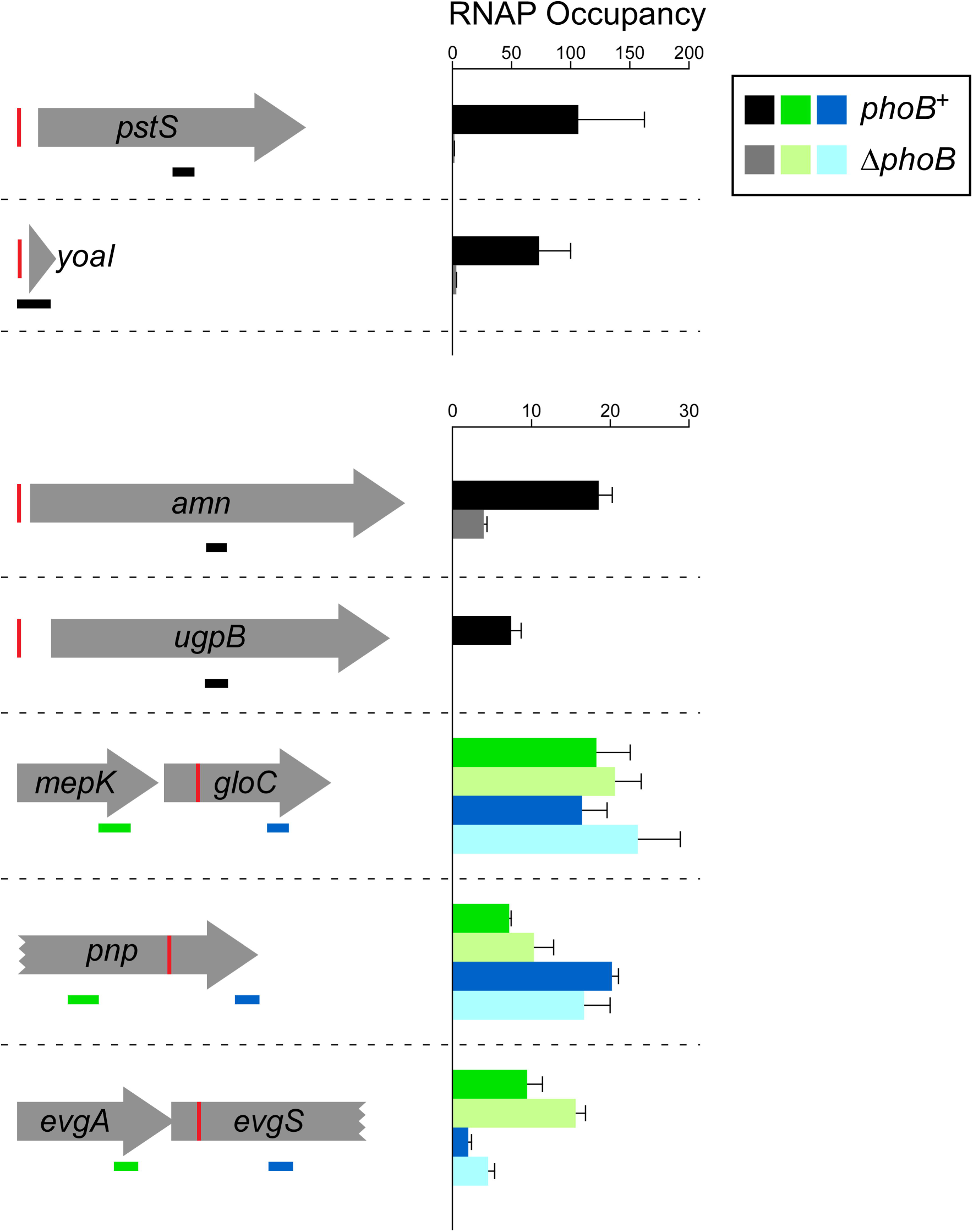
Differences in RNAP (β) occupancy in genes that are potential members of the *pho* regulon. RNAP (β) occupancy measured by ChIP-qPCR (see Methods for details of how occupancy units are calculated) in wild-type MG1655 (dark bars) and MG1655 Δ*phoB* (CDS091; light bars) for regions within genes that are potential members of the *pho* regulon. Schematics on the left show genes with upstream or internal PhoB sites (red vertical lines). The horizontal bars indicate the positions of PCR amplicons used in ChIP-qPCR; black bars indicate amplicons within genes that have upstream PhoB sites, green bars indicate amplicons upstream of intragenic PhoB sites, and blue bars indicate amplicons downstream of intragenic PhoB sites. Values represent the average of three independent biological replicates; error bars represent one standard deviation.

### PhoB-dependent recruitment of initiating RNAP

The majority of the PhoB binding sites identified by ChIP-seq were not associated with regulation detectable by RNA-seq. We hypothesized that this could be due to three reasons: (i) the binding sites are non-regulatory, (ii) regulation is condition-specific and/or requires additional factors, or (iii) PhoB regulates transcription of short, unstable, non-coding RNAs that are not detectable by conventional RNA-seq. To test the latter possibility, we used ChIP-seq to measure the association of σ^70^ in regions close to PhoB binding sites. σ^70^ is rapidly released from RNAP upon the transition from transcription initiation to elongation (44); thus, σ^70^ occupancy on DNA, as measured by ChIP-seq, is an indication of the level of association of initiating RNAP with DNA. Since transcription initiation occurs prior to RNA processing, σ^70^ occupancy can be observed even at promoters of highly unstable RNAs (45).

To measure the effects of PhoB on RNAP holoenzyme recruitment, we performed ChIP-seq of σ^70^ in wild-type and Δ*phoB* strains grown in low-phosphate medium. Normalized σ^70^ occupancy was calculated for 400 bp windows surrounding each PhoB binding site to systematically assess σ^70^binding at these sites (Figure 6). Three PhoB binding sites showed large reductions (>19-fold) in σ^70^occupancy in the Δ*phoB* strain relative to wild-type. Two of these sites are associated with the *phoB* gene itself; σ^70^ occupancy measurements at these sites are impacted by the loss of associated DNA sequence resulting from deletion of *phoB*. The third PhoB site is the regulatory site upstream of *pstS*. We conclude that PhoB activates *pstS* transcription at the level of RNAP recruitment, as suggested by structural models of the DNA:PhoB:RNAP complex (46–48). PhoB binding sites upstream of *yoaI*, *phoA*, *mglB*, *phnC*, and *phoH*, showed >2-fold lower σ^70^ occupancy in the Δ*phoB* strain relative to wild-type, suggesting that PhoB recruits initiating RNAP to these promoters. These data are largely consistent with the RNA-seq data showing >2-fold differential expression of *yoaI*, *phoA*, *phnC* and *yoaI* between wild-type and Δ*phoB* cells (Table 1). For most other PhoB sites, including almost all intragenic sites, σ^70^ occupancy was low in both wild-type and Δ*phoB* strains (Figure 6), strongly suggesting that these sites are not associated with active promoters under the growth conditions used. The remaining sites were associated with substantial σ^70^ occupancy that was similar in both wild-type and Δ*phoB* strains, suggesting that they are close to active promoters whose activity is independent of PhoB under the conditions tested.

**Figure 6.**
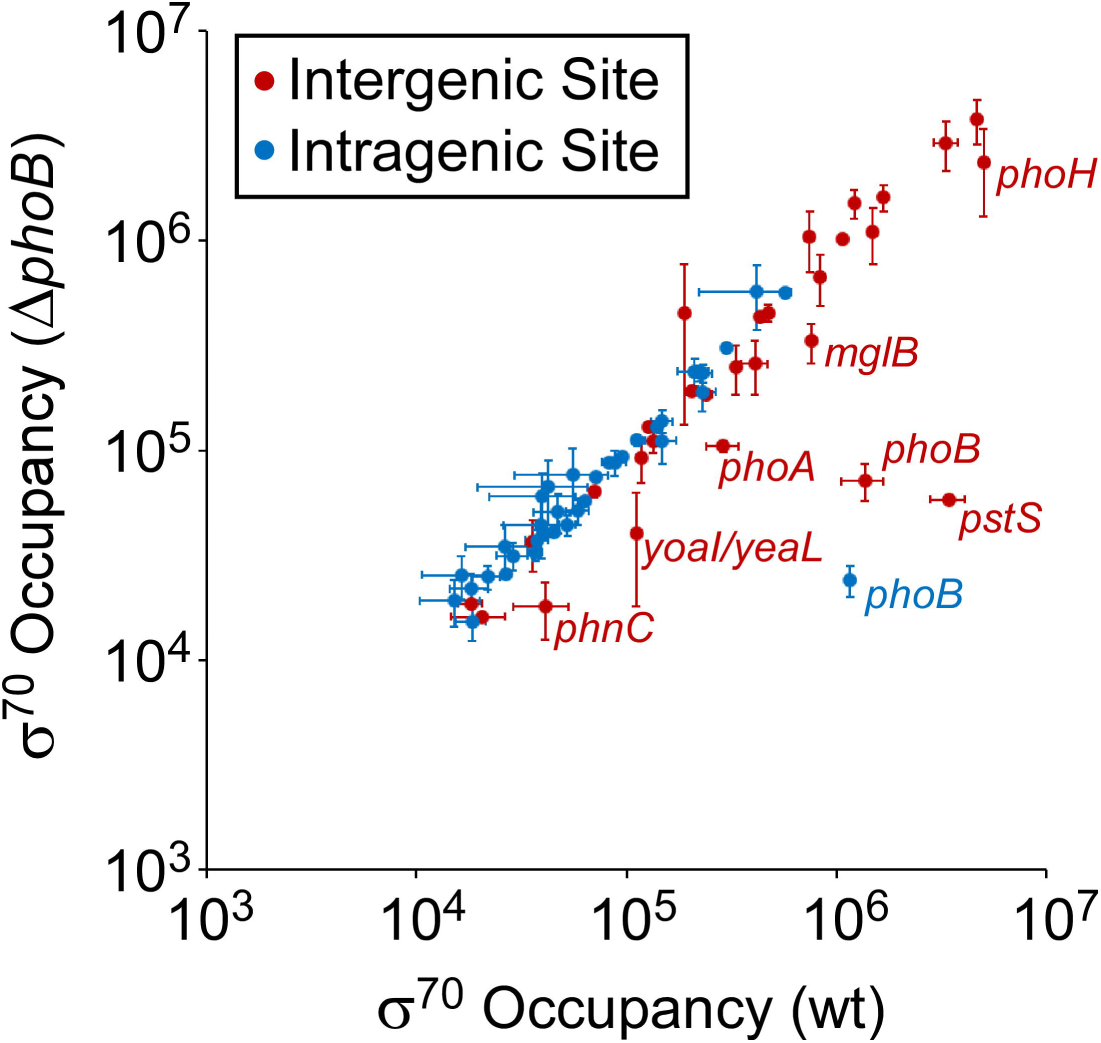
Differences in σ^70^ occupancy around PhoB binding sites between wild-type and Δ*phoB* cells. The scatter-plot shows normalized σ^70^ occupancy in wild-type MG1655 and MG1655 Δ*phoB* (DMF84) for the 400 bp regions surrounding PhoB binding sites identified by ChIP-seq. Each datapoint represents a PhoB binding site. Intergenic binding sites are indicated by red datapoints; intragenic binding sites by blue datapoints. Genes associated with PhoB binding sites are labeled with the gene name in cases where σ^70^ occupancy differs >2-fold between wild-type and Δ*phoB* cells. Values represent the average of two independent biological replicates; error bars represent +/- one standard deviation.

### H-NS co-associates with many intragenic PhoB sites, but does not block RNAP recruitment

We noted that 18 PhoB sites (12 intragenic and 6 intergenic), representing 28% of all sites identified by ChIP-seq, are in regions bound by the nucleoid-associated protein H-NS (49). Thus, PhoB sites are significantly enriched in H-NS-bound regions, which only represent 17% of the genome (Binomial test *p* = 0.02). Since H-NS is known to silence transcription (50), we hypothesized that the lack of detectable PhoB-dependent regulation at some sites may be due to the silencing effects of H-NS. To test this hypothesis, we repeated the σ^70^ ChIP-seq experiment in Δ*hns* and Δ*hns* Δ*phoB* strains. Comparison of σ^70^ occupancy between wild-type and Δ*hns* strains revealed substantially increased occupancy around some PhoB binding sites in the Δ*hns* strain, with most of these sites being intragenic (Figure 7A). Indeed, we observed widespread increases in σ^70^ occupancy at promoters genome-wide in the Δ*hns* strain relative to wild-type; most of the promoters showing increased σ^70^ association are located in regions of high H-NS occupancy (Figure 7B) (49). These data are consistent with our earlier study showing widespread transcriptional silencing by H-NS, particularly within genes (45). We next compared σ^70^ occupancy around PhoB binding sites between Δ*hns* and Δ*hns*Δ*phoB* strains (Figure 8). As for *hns*^+^ cells, the only large differences (>5-fold) in σ^70^ occupancy were associated with the PhoB sites at *phoB* and *pstS*. Interestingly, we did not observe differences >1.5-fold in σ^70^ occupancy at any other sites, including the sites upstream of *yoaI*, *phoA*, *mglB*, *phnC*, and *phoH*. However, the differences in σ^70^ occupancy we observed for *phoB* and *pstS* sites were between 2- and 4-fold lower than differences observed in the *hns*^+^ strains at the same sites. Hence, the more subtle differences in σ^70^ occupancy observed at the sites upstream of *yoaI*, *phoA*, *phnC*, and *phoH* in the *hns*^+^ strains may have escaped detection in the Δ*hns* strains. The lower effect of PhoB on σ^70^ occupancy in the Δ*hns* strain background may be due to the large-scale redeployment of RNAP that occurs in the absence of H-NS (51).

**Figure 7.**
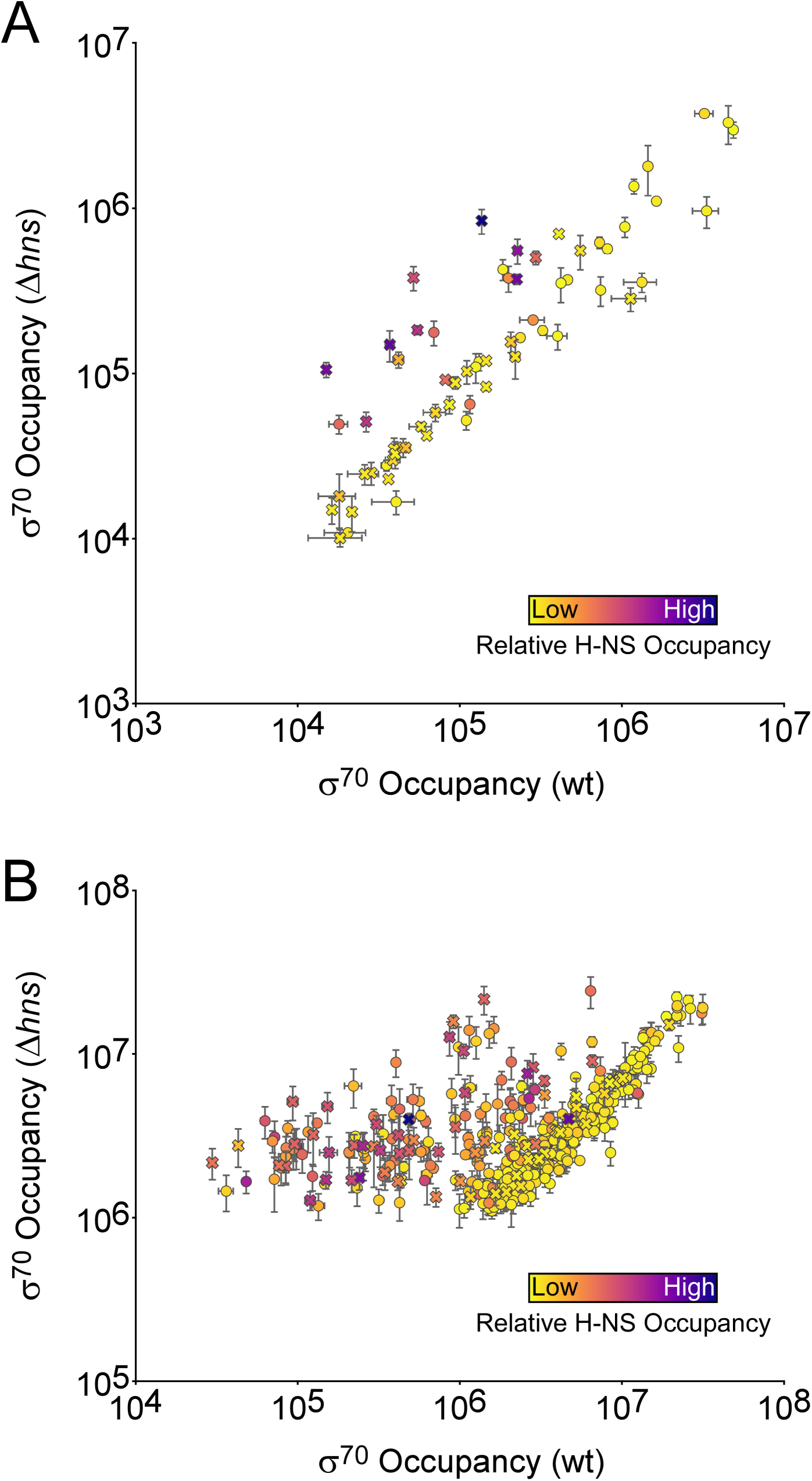
H-NS suppresses transcription from many promoters. **(A)** The scatter-plot shows normalized σ^70^ occupancy in wild-type MG1655 and MG1655 Δ*hns* (AMD565a) for the 400 bp regions surrounding PhoB binding sites identified by ChIP-seq. Each datapoint represents a PhoB binding site. The color of each datapoint indicates the level of H-NS occupancy at the corresponding site (49). Intragenic PhoB sites are represented by crosses; intergenic PhoB sites are represented by circles. **(B)** The scatter-plot shows normalized σ^70^ occupancy in wild-type MG1655 and MG1655 Δ*hns* (AMD565a) for all σ^70^ binding sites identified by ChIP-seq from MG1655 Δ*hns* (AMD565a) cells. The color of each datapoint indicates the level of H-NS occupancy at the corresponding site (49). Intragenic PhoB sites are represented by crosses; intergenic PhoB sites are represented by circles. For both (A) and (B), values represent the average of two independent biological replicates; error bars represent +/- one standard deviation.

**Figure 8.**
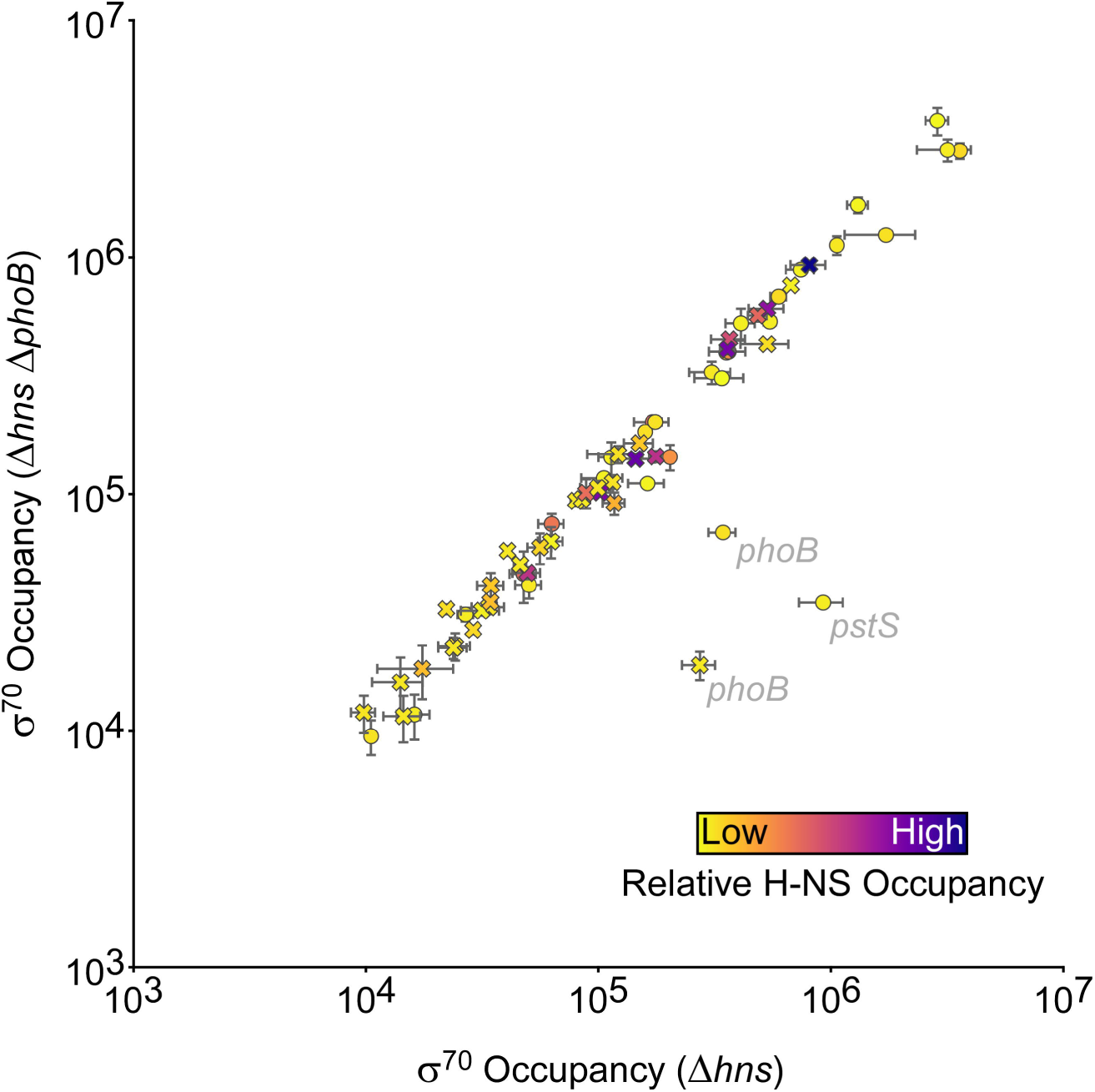
H-NS does not suppress PhoB-dependent effects on recruitment of initiating RNA polymerase. The scatter-plot shows normalized σ^70^ occupancy in wild-type MG1655 Δ*hns* (AMD565a) and MG1655 Δ*hns* Δ*phoB* (DMF85) for the 400 bp regions surrounding PhoB binding sites identified by ChIP-seq. Each datapoint represents a PhoB binding site. The color of each datapoint indicates the level of H-NS occupancy at the corresponding site (49). Intragenic PhoB sites are represented by crosses; intergenic PhoB sites are represented by circles. Genes associated with PhoB binding sites are indicated in cases where σ^70^ occupancy differs >2-fold between MG1655 Δ*hns* (AMD565a) and MG1655 Δ*hns* Δ*phoB* (DMF85) cells. Values represent the average of two independent biological replicates; error bars represent +/- one standard deviation.

While PhoB sites are enriched within H-NS-bound regions, H-NS does not appear to modulate PhoB activity at any site. We hypothesized that the enrichment of PhoB binding within H-NS-bound regions is due simply to the nucleotide content of the PhoB binding site; like H-NS-bound regions, the PhoB binding site has a higher A/T-content than the genome as a whole. To test this hypothesis, we scrambled the sequence of every PhoB binding site identified by ChIP-seq. We then derived a position weight matrix (PWM) from these scrambled sites and scored every genomic sequence for a match to this PWM. Strikingly, 36% of the top 1,000 scoring positions are within regions bound by H-NS (49). We conclude that the enrichment of PhoB binding sites within H-NS-bound regions is likely due to the A/T-rich nature of the binding motif.

### Sequence conservation of PhoB binding sites

Sequence conservation of a DNA binding site is often an indication that the site is functional (52). We determined the sequence conservation of the 59 *E. coli* PhoB sites identified by ChIP-seq for which we could identify an instance of the PhoB binding motif. Specifically, we scored homologous regions from 29 diverse γ- proteobacterial species for matches to the PhoB binding site motif (Figure 2C). The DNA-binding domain of PhoB is highly conserved across these species (Figure S1). As shown in Figure 9, the PhoB binding sites upstream of *pstS*, *phoB*, *phoA*, and *ugpB*, are broadly conserved. The PhoB binding sites upstream of *phoE* and *phoH* are conserved, albeit to a lesser degree. The PhoB binding site upstream of *phnC* is conserved in only a few species, suggesting that *phnC* is not a core member of the *pho* regulon. Among the novel PhoB binding sites, the best conserved is the site upstream of *rmf*, with strong matches to the PhoB DNA binding motif found upstream of *rmf* in most species analyzed. We detected PhoB upstream of *rmf* by ChIP-seq (Table 1) but did not detect significant PhoB-dependent regulation at the level of RNA abundance or RNAP recruitment. PhoB binding sites upstream of *agp*, *rpoH*, *cusR*/*cusC*, and *yoaI* were also conserved, albeit to a less degree, similar to sites upstream of *phoE* and *phoH*. The remaining intergenic PhoB sites are not well conserved, with most having few or no strong matches to the PhoB DNA binding motif outside of *E. coli*. Lastly, we examined conservation of intragenic PhoB sites. In most cases, these sites have little or no conservation; however, PhoB sites within *flhD*, *phoB*, and *pnp* are conserved among roughly half the species examined. This conservation may reflect a conserved function for the PhoB binding site, or could be due to sequence constraints on the codons.

**Figure 9.**
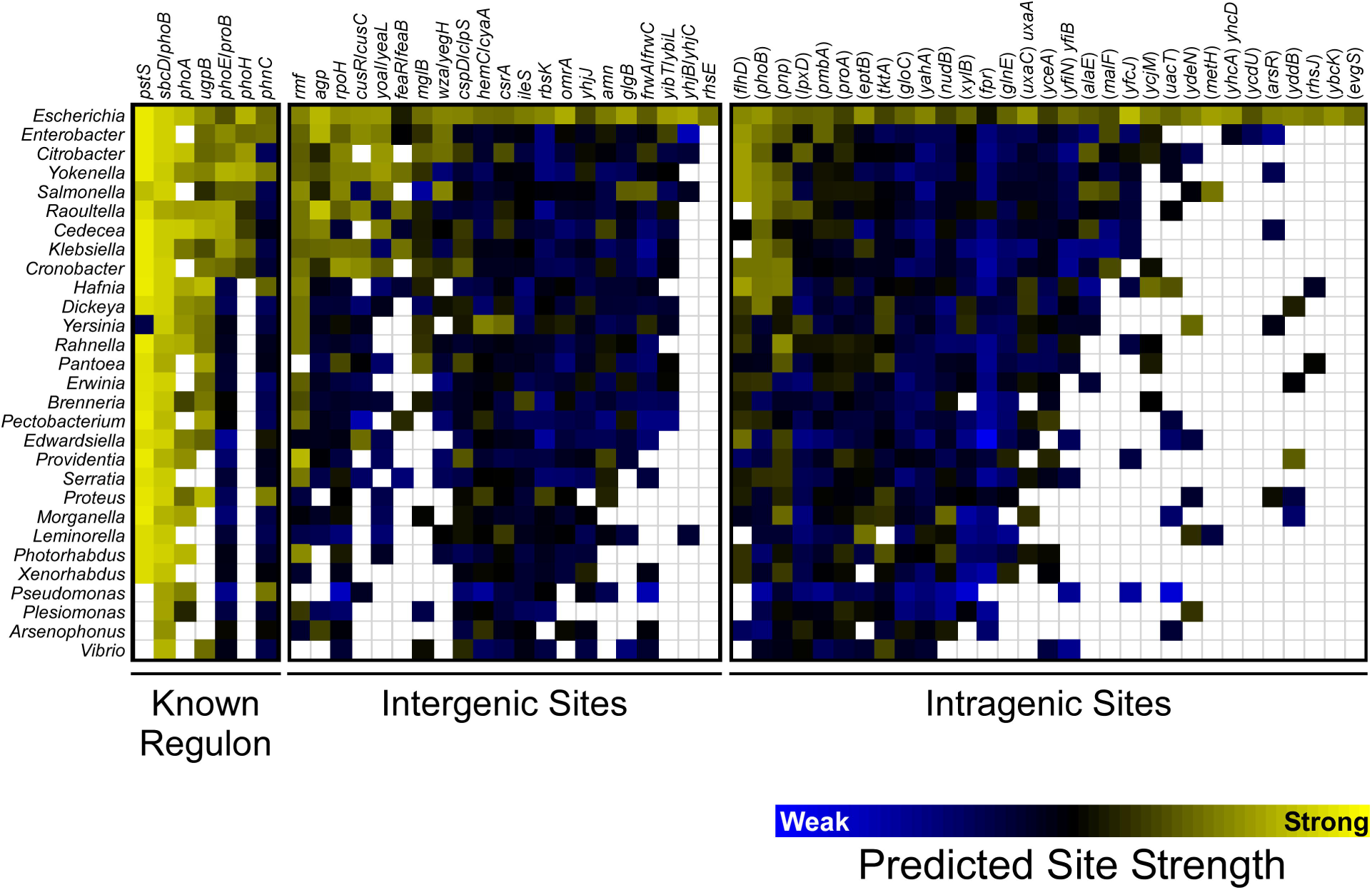
Conservation of PhoB binding sites across γ-proteobacterial species. Heat map showing conservation of PhoB binding sites across selected γ-proteobacterial species. Columns represent PhoB binding sites from *E. coli*, divided into known *pho* regulon binding sites, intergenic sites, and intragenic sites. The associated genes are indicated above each column. Rows represent species for different γ-proteobacterial genera, as indicated to the left of each row. The color of each square indicates the predicted strength of the best-scoring putative PhoB binding site in a region that is homologous to the corresponding region in *E. coli*. Binding site strength was predicted using a position weight matrix derived from the *E. coli* PhoB binding site motif (Figure 2B). The color scale is shown below the heat map, with yellow indicating stronger predicted binding site strength, and blue indicating weaker predicted binding site strength. White indicates the absence of a homologous region in the indicated species.

## DISCUSSION

### Comprehensive reassessment of the pho regulon in E. coli and beyond

By combining ChIP-seq and RNA-seq, we are able to reassess the *pho* regulon, with high resolution assignment of PhoB binding sites. As described above, the sensitivity and resolution of an earlier ChIP-chip study were substantially lower, precluding a comprehensive reassessment of the *pho* regulon (43). Previous studies disagree on which genes comprise the *pho* regulon in *E. coli*; however, most studies agree that the *pho* regulon includes the following operons: *pstSCAB*-*phoU*, *phoA*, *phoH*, *phnCDEFGHIJKLMNOP*, *phoBR*, *phoE*, *ytfK*, *ugpBAECQ*, and *psiE* (30, 33); *waaH* is considered in some studies to be in the *pho* regulon (33). Our data are largely consistent with these assignments, but provide strong evidence against *ytfK*, *psiE*, and *waaH* being members of the *pho* regulon. Specifically, we did not detect PhoB binding near any of these genes (Table 1), and we did not detect PhoB-dependent regulation of *psiE* (Table S1). It is formally possible that the C-terminal FLAG tags on PhoB altered its binding specificity or reduced its affinity for DNA such that we failed to detect binding to sites upstream of *ytfK*, *psiE*, and *waaH*. Nonetheless, in such a scenario, these binding sites would presumably have relatively low affinity for PhoB.

Our data also rule out several other putative *pho* regulon genes: *asr*, *eda*, *argP*, and *pitB* that were not associated with detectable PhoB binding or PhoB-dependent regulation (40, 53–57). Similarly, we did not detect binding of PhoB upstream of the sRNA-encoding gene *esrL*, despite a recent report of PhoB binding to this region in enteropathogenic *E. coli*, with the sequence of the reported PhoB site being identical in *E. coli* K- 12 (58). By contrast, our ChIP-seq data support the assignment of *amn* and *yoaI* as *pho* regulon members, as has previously been suggested based on limited experimental evidence (40–42). Our ChIP-seq and RNA-seq data identify novel *pho* regulon members with confidence: *cusC*, *feaR*, *yibT*, and *agp*. These genes all have PhoB binding sites upstream, and were significantly, differentially expressed >2-fold between wild-type and Δ*phoB* strains. We note that direct positive regulation of *cusC* and direct negative regulation of *feaR* by PhoB have been suggested previously (43). Our data provide no evidence to suggest that there are unannotated transcripts regulated by PhoB, or transcripts whose regulation by PhoB is masked by H-NS (Figures 6 + 8). Lastly, our data do not support direct regulation by PhoB sites located within genes (Figure 5).

Phylogenetic analysis of PhoB binding sites highlights a highly conserved set of *pho* regulon genes within the γ-proteobacteria: *pstS*, *phoB*, *phoA*, *ugpB*, and associated operonic genes (Figure 9). Consistent with this, direct PhoB regulation of the *pstS* and *phoB* transcripts has been described for the more distantly related α- proteobacterium *Caulobacter crescentus* (59). *phoE*, *phoH*, and *yoaI* represent a second set of conserved *pho* regulon genes, although their conservation is more phylogenetically restricted. Interestingly, while PhoB regulation of *phnC* and associated operonic genes does not appear to be widely conserved, we did observe evidence for strong PhoB sites upstream of *phnC* in a small set of species, and *phnC* is known to be a direct regulatory target of PhoB in *C. crescentus* (59), suggesting that *phnC* may have a niche-specific function in phosphate homeostasis.

The phylogenetic pattern of PhoB binding site conservation for sites upstream of *rmf, agp*, *rpoH*, and *cusR*/*cusC* suggests these genes may be part of the conserved *pho* regulon. We observed significant differential expression of *cusC* and *agp* between wild-type and Δ*phoB* cells. By contrast, we did not observe significant differential expression of *rmf* or *rpoH*. We speculate that regulation of *rmf* and *rpoH* by PhoB is integrated with regulation by other transcription factors, such that PhoB-dependent changes in expression are only detectable under specific growth conditions. Consistent with this idea, transcription of *rmf* has been shown to be regulated by ppGpp (60, 61) and CRP (62), and possibly by additional transcription factors (63) and diverse stress conditions (64). PhoB binding sites upstream of *agp*, *rpoH*, and *cusR*/*cusC* are conserved in largely the same set of species as binding sites upstream of *phoE*, *phoH*, and *yoaI*, suggesting that these species share a common set of *pho* regulon genes.

### Most intragenic PhoB sites appear to be non-functional, and are not under selective pressure

Our data argue against intragenic PhoB sites having regulatory activities of the types that have been described previously for intragenic TF sites; specifically, regulation of transcription from an intragenic promoter (5, 7–12), or regulation of the overlapping gene either by roadblock repression or a novel mechanism (3, 15–18). Indeed, most intragenic PhoB sites are associated with little or no local σ^70^ binding (Figure 6), indicating that PhoB binding alone is insufficient to recruit RNAP. Thus, it is likely that RNAP:σ^70^-interacting promoter elements are also necessary for PhoB-dependent recruitment of RNAP. Moreover, the spacing between PhoB sites and core promoter elements is likely to be important in determining whether PhoB recruits RNAP, since even intragenic PhoB sites that are close to intragenic promoters (i.e., those associated with high ChIP-seq signal for σ^70^) show no difference in σ^70^occupancy upon deletion of *phoB*. Consistent with this idea, structural models of the PhoB:RNAP:DNA complex formed at PhoB-activated promoters support strict spacing requirements between the *pho* box and core promoter elements (46–48, 65).

Widespread intragenic TF binding is emerging as a common phenomenon as more TFs are mapped using ChIP- seq (2–6, 66). Similarly, many σ factors have been shown to bind and initiate transcription from large numbers of intragenic promoters (8, 19, 20, 67–70). In the majority of cases tested, these intragenic binding sites are poorly conserved (69, 71), as is the case for intragenic PhoB sites (Figure 9). Based on the lack of detectable transcriptional activity, and the limited conservation of intragenic PhoB sites, we speculate that intragenic TF sites often arise due to genetic drift or selective pressures on overlapping sequences such as codons. Consistent with this idea, intragenic PhoB sites tend to be weaker (lower ChIP-seq enrichment) than intergenic sites (Mann-Whitney U test, *p* = 0.005). A previous study showed that the predicted number of intragenic binding sites for many bacterial TFs is the same in actual genome sequences as it is in randomized genome sequences, suggesting that intragenic TF binding sites are common and arise largely due to genetic drift (28). We further speculate that the fitness cost of intragenic PhoB sites is low. Intragenic TF binding sites can therefore be considered genomic “noise”. Interestingly, the vast majority of PhoB binding events detected in *C. crescentus* are intergenic (59), suggesting that intragenic PhoB binding in *C. crescentus* may be associated with a fitness cost. Finally, we cannot rule out the possibility that intragenic PhoB sites in *E. coli* are functional. For example, they could contribute, *en masse*, to titration of PhoB, they could facilitate DNA looping that impacts chromosome structure, as has been suggested for some TFs (22–27), or they could regulate transcription by unknown mechanisms.

We have comprehensively mapped the PhoB regulon by assessing PhoB binding, PhoB-dependent transcriptome changes, and PhoB-dependent RNAP recruitment. We identified novel *pho* regulon members, some of which are modestly conserved across other genera, and identified many seemingly non-functional PhoB binding sites inside genes. We conclude that a combination of binding site information (e.g., ChIP-seq) and regulatory information (e.g., RNA-seq) is required to accurately define the regulons of most TFs.

## MATERIALS AND METHODS

### Strains and plasmid

*E. coli* MG1655 and its derivatives were used for this study. The strains and plasmid used are listed in Table 2. All oligonucleotides used are listed in Table S2. For ChIP-seq, PhoB was C-terminally epitope tagged with a 3x-FLAG tag. The tag was inserted at the native *phoB* locus using FRUIT recombineering (72) using oligonucleotides JW2973 and JW2974. MG1655 Δ*phoB* (DMF84) and Δ*hns* (AMD565a) strains were constructed by P1 transduction from Keio collection strains (73) into MG1655. The *kan^R^* genes were removed by FLP-recombinase expressed from pCP20, as previously described (74). The MG1655 Δ*hns* Δ*phoB* strain was made in the same manner, with deletions introduced sequentially. MG1655 Δ*phoB* (CDS091) was constructed using FRUIT (72) using oligonucleotides JW6280, JW6281, JW6294 and JW6295. Note that there are two MG1655 Δ*phoB* strains used in this study. DMF34 was used for σ^70^ ChIP-seq experiments, and CDS091 was used for qRT-PCR, RNA-seq, and RNAP ChIP-qPCR experiments.

**Table 2.**
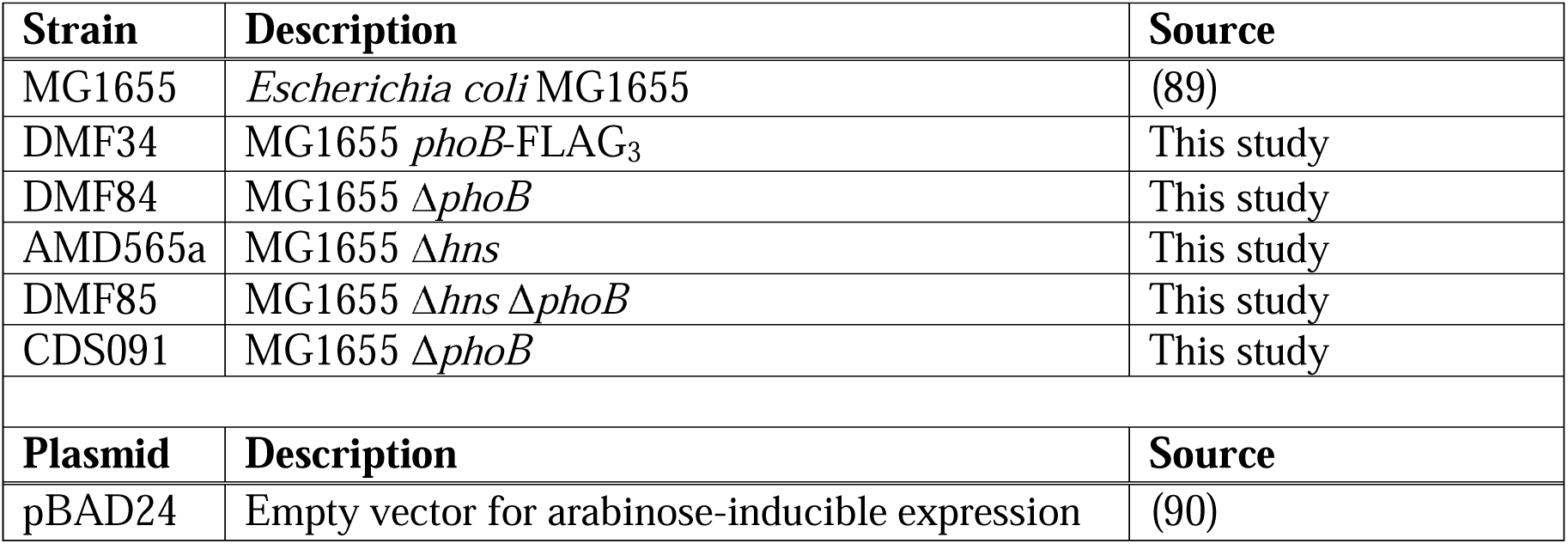
List of strains and plasmid used in this study.

### Quantitative reverse transcription PCR

Quantitative RT-PCR was performed based on a previous study (69). MG1655 + pBAD24, CDS091 (MG1655 Δ*phoB*) + pBAD24, or DMF34 + pBAD24 cells were grown at 37° C with aeration in MOPS minimal medium with 0.2 mM K_2_PO_4_, 0.4% glucose and 100 µg/ml ampicillin to an OD_600_ of 0.5-0.6. Arabinose was added to a final concentration of 0.2% for 7 minutes or 20 minutes, with one or two of the three replicate samples for each strain receiving arabinose for 7 minutes. Thus, the replicate samples were not always consistent with respect to the extent of growth after arabinose addition. However, since arabinose is not expected to affect *pstS* expression, all samples for a single strain were treated as replicates regardless of whether cells were grown for 7 minutes or 20 minutes after arabinose addition. RNA was prepared as described for RNA-seq. RNA was reverse-transcribed using SuperScript III reverse transcriptase (Invitrogen) according to the manufacturer’s instructions. A control reaction omitting reverse transcriptase was performed. 1% of the cDNA (or negative control) was used as a template in a quantitative real time PCR using an Applied Biosystems 7500 Fast real time PCR machine, with primers JW156 + JW157 for amplifying the *minD* control gene, and JW7802 + JW7803 for amplifying *pstS*. Relative expression of *pstS* was determined by the ΔC_T_ method, normalizing to *minD* expression.

### ChIP-seq

For low phosphate growth experiments, cells were grown at 37° C with aeration in MOPS minimal medium with 0.2 mM K_2_HPO_4_ and 0.4% glucose, as previously described (37, 75). For high phosphate growth experiments, cells were grown in MOPS minimal medium with 1.32 mM K_2_HPO_4_ and 0.4% glucose. Subcultures were inoculated 1:100, and grown at 37° C with aeration to an OD_600_ of 0.5-0.7. ChIP-seq libraries were prepared as previously described (76). Libraries were prepared from two biological replicate cultures for each experimental group. For DMF34 (MG1655 phoB-FLAG_3_) or MG1655, PhoB-FLAG_3_ (or the untagged control) was immunoprecipitated with 2 μL of α-FLAG M2 monoclonal antibody (Sigma). For MG1655, DMF84 (Δ*phoB*), AMD565a (MG1655 Δ*hns*), and DMF85 (MG1655 Δ*phoB* Δ*hns*), σ^70^ was immunoprecipitated with 1 μL of α-σ^70^ antibody (Neoclone). Libraries were sequenced on a HiSeq 2000 (Illumina) by the University at Buffalo Next-Generation Sequencing Core Facility or a NextSeq (Illumina) by the Wadsworth Center Applied Genomic Technologies Core Facility.

### Analysis of PhoB-FLAG_3_ ChIP-seq Data

Duplicate ChIP-seq data for (i) MG1655 PhoB-FLAG_3_ (DMF34) grown in low phosphate conditions, (ii) untagged MG1655 grown in low phosphate conditions, and (iii) MG1655 PhoB-FLAG_3_ (DMF34) grown in high phosphate conditions, were aligned to the *E. coli* MG1655 genome (NC_000913.3) using CLC Genomics Workbench (version 8). ChIP-seq peaks were called using a previously described analysis pipeline (8).

### Analysis of RNAP occupancy around PhoB binding sites

Wild-type MG1655 and MG1655 Δ*phoB* (CDS091) cells were grown at 37° C with aeration to an OD_600_ of 0.6- 0.7 in MOPS minimal medium with 0.2 mM K_2_HPO_4_ and 0.4% glucose, as previously described (37, 75). ChIP was performed as previously described (76), using 1 µl anti-β (RNA polymerase subunit) antibody (BioLegend catalog #663903). ChIP and input samples were analyzed using an ABI 7500 Fast real-time PCR machine. Enrichment of ChIP samples was calculated relative to a control region within *bglB*, which is transcriptionally silent. RNAP Occupancy Units represent background-subtracted fold–enrichment over the control region. Oligonucleotides used for qPCR were JW125 + JW126 (*bglB*), JW10937 + JW10938 (*yoaI*), JW10939 + JW10940 (*amn*), JW10941 + JW10942 (*pstS*), JW10943 + JW10944 (*ugpB*), JW10945 + JW10946 (*mepK*), JW10947 + JW10948 (*gloC*), JW10949 + JW10950 (*evgA*), JW10951 + JW10952 (*evgS*), JW10953 + JW10954 (*pnp* upstream), JW10955 + JW10956 (*pnp* downstream). For statistical analysis of relative changes in RNAP occupancy upstream and downstream of intragenic PhoB sites, we used the mean and standard deviation values for ChIP-qPCR occupancy to generate 1,000 simulations for occupancy at each region tested. We then determined how frequently the ratio of predicted RNAP occupancy downstream:upstream of an intragenic PhoB site was higher in Δ*phoB* cells than wild-type cells. We repeated this simulation 10 times to estimate a *p*-value for each of the three intragenic PhoB sites tested.

### Analysis of ^70^ occupancy around PhoB binding sites

Duplicate ChIP-seq data for σ^70^ from wild-type MG1655, MG1655 Δ*phoB* (DMF84), MG1655 Δ*hns* (AMD565a), and MG1655 Δ*hns* Δ*phoB* (DMF85) were aligned to the *E. coli* MG1655 genome (NC_000913.3) using Rockhopper (version 2.03, default parameters) (77), which also calculated the depth of sequence coverage at all genomic positions on each strand, normalized for total sequence read count. A custom Python script was used to determine the relative sequence read coverage from each σ^70^ ChIP-seq dataset in 400 bp windows centered on each PhoB ChIP-seq peak (coordinates listed in Table 1).

### Analysis of σ^70^ occupancy at promoters in wild-type and Δhns strains

Duplicate ChIP-seq data for σ^70^ for wild-type MG1655 or MG1655 Δ*hns* (AMD565a) were aligned to the *E. coli* MG1655 genome (NC_000913.3) using CLC Genomics Workbench (version 8). ChIP-seq peaks were called using a previously described analysis pipeline (8). A custom Python script was used to determine the relative sequence read coverage from each σ^70^ChIP-seq dataset in 50 bp windows centered on each σ^70^ ChIP- seq peak from MG1655 Δ*hns* (AMD565a).

### Determining H-NS occupancy from published ChIP-seq data

H-NS ChIP-seq occupancy was determined from published data (49). Specifically, genome coordinates for σ^70^ ChIP-seq peaks were converted from NCBI genome sequence version U00096.3 to U00096.2 at https://biocyc.org/ECOLI/map-seq-coords-form?chromosome=COLI-K12. H-NS occupancy was determined as the average from four normalized sequence read coverage files available from EBI ArrayExpress, accession number E-MTAB-332.

### RNA-seq

MG1655 + pBAD24 or CDS091 (MG1655 Δ*phoB*) + pBAD24 cells were grown at 37° C with aeration in MOPS minimal medium with 0.2 mM K_2_PO_4_, 0.4% glucose and 100 µg/ml ampicillin to an OD_600_ of 0.5-0.6 Arabinose was added to a final concentration of 0.2% for 7 minutes. Note that addition of arabinose is not expected to impact expression of PhoB-regulated genes. RNA was isolated using a modified hot-phenol method, as previously described (76). Samples were treated with Turbo DNase (Ambion) to remove genomic DNA, ribosomal RNA was removed using the Ribo-Zero rRNA removal kit for Gram-negative bacteria (Epicentre/Illumina), and libraries were prepared with the ScriptSeq Complete kit for bacteria (Epicentre/Illumina) (76). Libraries were sequenced on a HiSeq 2000 (Illumina) by the University at Buffalo Next-Generation Sequencing Core Facility. RNA-seq data were aligned to the *E. coli* MG1655 genome (NC_000913.3) using BWA for Illumina (v0.5.9-r16) (78) on Galaxy (usegalaxy.org) (79). Read counting, normalization, and differential expression analysis were performed in R using GenomicAlignments (v1.28) *summarizeOverlaps* (80) and DEseq2 (v1.32, betaPrior = FALSE) (81).

### PhoB Motif Discovery and Analysis

100 bp sequences surrounding PhoB ChIP-seq peaks were extracted and analyzed using MEME (version 5.1.0, default parameters) (82, 83). The position of the inferred motif relative to ChIP-seq peak centers was analyzed using Centrimo (version 5.1.0, default parameters) (84) through the MEME-ChIP tool (85).

To determine whether the nucleotide content of the PhoB binding site motif contributes to the association of PhoB binding sites with H-NS-bound regions, we first scrambled each PhoB binding site individually using a custom Python script. We then compiled the scrambled sites into a PWM and searched the *E. coli* MG1655 genome (NC_000913.3) for the top 1000 matches to this PWM using FIMO (version 5.1.0, default parameters) (86).

### Analysis of PhoB binding site conservation

Binding site conservation analysis was performed as described previously (71). Protein sequences were aligned using Clustal Omega (87) and visualized using MView (88). The genomes analyzed were *Arsenophonus nasoniae* DSM 15247, *Brenneria sp*. EniD312, *Cedecea davisae* DSM 4568, *Citrobacter rodentium* ICC168, *Cronobacter sakazakii* ATCC BAA-894, *Dickeya dadantii* 3937, *Edwardsiella tarda* EIB202, *Enterobacter cloacae* subsp. *cloacae* ATCC 13047, *Erwinia amylovora* ATCC 49946, *Escherichia coli* str. K-12 substr. MG1655, *Hafnia alvei* ATCC 51873, *Klebsiella pneumoniae* KCTC 2242, *Leminorella grimontii* ATCC 33999 = DSM 5078, *Morganella morganii* subsp. *morganii* KT, *Pantoea agglomerans* 299R, *Pectobacterium atrosepticum* SCRI1043, *Photorhabdus asymbiotica*, *Plesiomonas shigelloides* 302-73, *Proteus mirabilis* HI4320, *Providencia stuartii* MRSN 2154, *Pseudomonas aeruginosa* PAO1, *Rahnella sp*. Y9602, *Raoultella ornithinolytica* B6, *Salmonella enterica* str P125109, *Serratia marcescens* FGI94, *Vibrio cholerae* M66-2, *Xenorhabdus bovienii* SS-2004, *Yersinia pestis* KIM 10, and *Yokenella regensburgei* ATCC 43003.

## ACKNOWLEDGEMENTS

We thank the University at Buffalo Next-Generation Sequencing Core Facility and the Wadsworth Center Applied Genomic Technologies Core Facility for DNA sequencing. We thank the Wadsworth Center Tissue Culture and Media Core Facility and Glassware Facility for technical support. This material is based on work supported by the National Science Foundation Graduate Research Fellowship under grant number DGE- 1060277 (DMF). DMF was also supported by National Institutes of Health training grant T32AI055429. This work was also supported by a National Institutes of Health Director’s New Innovator Award DP2OD007188 (JTW) and National Institutes of Health grants R01GM114812 and R35GM144328 (JTW).

## ACCESSION NUMBERS

ChIP-seq data are available at EBI ArrayExpress using accession number E-MTAB-9293. RNA-seq data are available at EBI ArrayExpress using accession number E-MTAB-9591.

## SUPPLEMENTARY TABLES

**Table S1. RNA-seq analysis.**

**Table S2. List of oligonucleotides used in this study.**

