## Supplementary material for "Genome-wide mapping of the *Escherichia coli* PhoB regulon reveals many transcriptionally inert, intragenic binding sites": Figure S1

| | cov | id | |
|---|---|---|---|
| *Escherichia* | 100.0% | 100.0% | ----------MARRILVVEDEAPIREMVCFVLEQNGFQPVEAEDYDSAVNQLNEPWPDLILLDWMLPGGSGIQFIKHLKR |
| *Enterobacter* | 100.0% | 96.5% | ----------MARRILVVEDEAPIREMVCFVLEQNGFQPVEAEDYDSAVNQLNEPWPDLILLDWMLPGGSGLQFIKHLKR |
| *Citrobacter* | 100.0% | 95.6% | ----------MARRILVVEDEAAIREMVCFVLEQNGFQPVEAEDYDTAVNQLNEPWPDLILLDWMLPGGSGLQFIKHLKR |
| *Yokenella* | 100.0% | 95.6% | ----------MARRILVVEDEAPIREMVCFVLEQNGFQPVEAEDYDSAVNQLNEPWPDLILLDWMLPGGSGLQFIKHIKR |
| *Salmonella* | 100.0% | 95.6% | ----------MARRILVVEDEAPIREMVCFVLEQNGFQPVEAEDYDSAVNKLNEPWPDLILLDWMLPGGSGLQFIKHLKR |
| *Raoultella* | 100.0% | 95.6% | ----------MARRILVVEDEAPIREMVCFVLEQNGFQPVEAEDYDSAVNQLNEPWPDLILLDWMLPGGSGLQFIKLLKR |
| *Klebsiella* | 100.0% | 95.2% | ----------MARRILVVEDEAPIREMVCFVLEQNGFQPVEAEDYDSAVNQLNEPWPDLILLDWMLPGGSGLQFIKLLKR |
| *Cronobacter* | 100.0% | 94.8% | ----------MARRILVVEDEAPIREMVCFVLEQNGFQPVEAEDYDSAVNQLNEPWPDLILLDWMLPGGSGIQFIKHIKR |
| *Cedecea* | 100.0% | 92.1% | ----------MARRILVVEDEAPIREMVSFVLEQNGFQSVEAEDYDSAVNLLIEPFPDLILLDWMLPGGSGIQFIKHLKR |
| *Hafnia* | 100.0% | 91.7% | ----------MARRILVVEDEAPIREMVCFVLEQNGYQPVEAEDYDSAVNSLSEPYPDLVLLDWMLPGGSGIQFIKHMKR |
| *Serratia* | 100.0% | 91.3% | ----------MARRILVVEDEAPIREMVCFVLEQNGYQPLEAEDYDSAVTRLSEPFPDLVLLDWMLPGGSGIQFIKHMKR |
| *Pantoea* | 100.0% | 90.4% | ----------MAKRILVVEDEAPIREMLCFVLEQNDYQPIEAEDYDSAVGKLIEPWPDLILLDWMLPGGSGIQFIKHLKR |
| *Brenneria* | 100.0% | 89.5% | ----------MARRILVVEDEAPIREMVCFVLEQNGYQPVEAEDYDSAVTQLSEPFPELVLLDWMLPGGSGLQFIKHMKR |
| *Dickeya* | 100.0% | 89.1% | ----------MARRILVVEDEAPIREMVCFVLEQNGYQPVEAEDYDSAVTRLAEPYPELVLLDWMLPGGSGLQFIKHMKR |
| *Rahnella* | 100.0% | 89.1% | ----------MARRILVVEDEAPIREMVCFVLEQNGYQAVEAEDFDSAIGQLVEPYPELILLDWMLPGGSGIQFIKHLKR |
| *Pectobacterium* | 100.0% | 89.1% | ----------MAKRILVVEDEAPIREMVCFVLEQNGYQPVEAEDYDSAVTQLSEPFPELVLLDWMLPGGSGLQFIKHMKR |
| *Erwinia* | 100.0% | 87.8% | ----------MAKRILVVEDEAPIREMLCFVLEQNDYQPIEAADYDSALSLLIEPWPDLILLDWMLPGGSGIQFIKHLKR |
| *Yersinia* | 100.0% | 86.2% | MTANILAGLMMARRILVVEDEAPIREMVCFVLEQNGYQPLEAEDYDSAVARLSEPFPDLVLLDWMLPGGSGIQFIKHMKR |
| *Edwardsiella* | 100.0% | 85.2% | ----------MSIRILVVEDETPIRDMVSFVLEQNGYQPLEAESYDGALSQLCEPYPDLILLDWMLPGGSGIQLIKQLKR |
| *Plesiomonas* | 100.0% | 84.3% | ----------MARRILVVEDEAPIRDMVCFVLEQKGYEPVEAEDYDAALSKMAEPYPDLILLDWMLPGGTGIQLIKHLKR |
| *Providencia* | 100.0% | 81.7% | ----------MARRILVVEDEAPIREMVCFVLEQNGFQSIEADDYDSAIAQLVDPLPDLVLLDWMIPGGSGIQVIKHMKR |
| *Xenorhabdus* | 100.0% | 81.7% | ----------MAKRILVVEDEVQIREMVCLVLEQNGYQTVEAEDYDVAVWRLSEPYPDLVLLDWMLPGGSGIQLIKQMKR |
| *Photorhabdus* | 100.0% | 81.2% | ----------MTRRILVVEDETPIREMVCFVLEKNGYQPVEAEDYDSALACLSEPYPDLVLLDWMIPGGSGLQIIKQMKR |
| *Morganella* | 100.0% | 81.2% | ----------MARRILVVEDEAPIREMVCFVLEQNGYQPIEADDYDAAIARLVEPFPDLVLLDWMIPGGSGIQVIKHMKR |
| *Vibrio* | 100.0% | 80.3% | ----------MSRRILVVEDEAPIREMLCFVLEQKGYQAVEAEDYDSAMSKLAEPFPDLVLLDWMLPGGSGINLIKHMKR |
| *Proteus* | 100.0% | 79.0% | ----------MARRILVVEDETAIREMICFVLEQNGFQPIEAEDYDTALSFLIDPYPDLVLLDWMIPGGSGIQVIKQMKR |
| *Leminorella* | 100.0% | 75.1% | ----------MMKRILIVEDEAPIREMISLVLEQHDYQTVEAGDLASAQAQLKEPYPDLVLLDWMLPGGSGIQFIKSMKR |
| *Arsenophonus* | 100.0% | 74.8% | -----MIGSIMVRRILVVEDETAIREMVCFVLEQNGFQTVEADDYDSAIAQLIEPYPALILLDWMIPGGSGIQLIAHMKR |
| *Pseudomonas* | 99.6% | 61.3% | ---------MVGKTILIVDDEAPIREMIAVALEMAGYECLEAENTQQAHAVIVDRKPDLILLDWMLPGTSGIELARRLKR |

| | cov | id | |
|---|---|---|---|
| *Escherichia* | 100.0% | 100.0% | ESMTRDIPVVMLTARGEEEDRVRGLETGADDYITKPFSPKELVARIKAVMRRISPMAVEEVIEMQGLSLDPTSHRVMAGE |
| *Enterobacter* | 100.0% | 96.5% | EAMTRDIPVVMLTARGEEEDRVRGLETGADDYITKPFSPKELVARIKAVMRRISPMAVEEVIEMQGLSLDPTSHRVMTGE |
| *Citrobacter* | 100.0% | 95.6% | EAMTRDIPVVMLTARGEEEDRVRGLETGADDYITKPFSPKELVARIKAVMRRISPMAVEEVIEMQGLSLDPTSHRVMTGD |
| *Yokenella* | 100.0% | 95.6% | EAMTRDIPVMMLTARGEEEDRVRGLETGADDYITKPFSPKELVARIKAVMRRISPMAVEEVIEMQGLSLDPTSHRVMTGE |
| *Salmonella* | 100.0% | 95.6% | EAMTRDIPVVMLTARGEEEDRVRGLETGADDYITKPFSPKELVARIKAVMRRISPMAVEEVIEMQGLSLDPGSHRVMTGD |
| *Raoultella* | 100.0% | 95.6% | EAMTRDIPVVMLTARGEEEDRVRGLETGADDYITKPFSPKELVARIKAVMRRISPMAVEEVIEMQGLSLDPSSHRVMTGE |
| *Klebsiella* | 100.0% | 95.2% | EAMTRDIPVVMLTARGEEEDRVRGLETGADDYITKPFSPKELVARIKAVMRRISPMAVEEVIEMQGLSLDPSSHRVMTGD |
| *Cronobacter* | 100.0% | 94.8% | EALTREIPVVMLTARGEEEDRVRGLETGADDYITKPFSPKELVARIKAVMRRISPMAVDEVIEMQGLSLDPSSHRVMTGE |
| *Cedecea* | 100.0% | 92.1% | EALTREIPVMMLTARGEEEDRVRGLEVGADDYITKPFSPKELVARIKAVMRRISPMAVEEVIEMQGLSLDPSSHRVMTGE |
| *Hafnia* | 100.0% | 91.7% | EALTREIPVMMLTARGEEEDRVRGLEVGADDYITKPFSPKELVARIKAVMRRISPMAVEEVIEVQGLSLDPTSHRVMANE |
| *Serratia* | 100.0% | 91.3% | EALTRDIPVMMLTARGEEEDRVRGLEVGADDYITKPFSPKELVARIKAVMRRISPMAVEEVIEMQGLSLDPSSHRVMANE |
| *Pantoea* | 100.0% | 90.4% | EAMTRDIPVMMLTARGEEEDRVRGLEVGADDYITKPFSPKELMARIKAVMRRISPMAVEEVIEMQGLSLDPSSHRVMSET |
| *Brenneria* | 100.0% | 89.5% | EALTRDIPVMMLTARGEEEDRVRGLEVGADDYITKPFSPKELVARIKAVMRRISPMAVEEVIEMRGLSLDPSSHRVTTEE |
| *Dickeya* | 100.0% | 89.1% | EALTRDIPVMMLTARGEEEDRVRGLEVGADDYITKPFSPKELVARIKAVMRRISPMAVEEVIEMRGLSLDPSSHRVTTEE |
| *Rahnella* | 100.0% | 89.1% | EALTREIPVMMLTARGEEEDRVRGLEVGADDYITKPFSPKELVARIKAVMRRISPMAVEEVIDMQGLSLDPSSHRVMANE |
| *Pectobacterium* | 100.0% | 89.1% | EALTRDIPVMMLTARGEEEDRVRGLEVGADDYITKPFSPKELVARIKAVMRRISPMAVEEVIEMRGLSLDPSSHRVTTEE |
| *Erwinia* | 100.0% | 87.8% | EAMTRDIPVMMLTARGEEEDRVRGLEVGADDYITKPFSPKELVARIKAVMRRIAPMAVEEVIEIQGLTLDPSSHRVMSGT |
| *Yersinia* | 100.0% | 86.2% | EALTRDIPVMMLTARGEEEDRVRGLEVGADDYITKPFSPKELVARIKAVMRRISPMAVEEVIEMQGLSLDPSSHRVMAND |
| *Edwardsiella* | 100.0% | 85.2% | EPATREIPVMMLTARGEEEDRVRGLEVGADDYITKPFSPKELVARIKAVMRRISPMALEETINLQGLSLDPVSHRVTAQD |
| *Plesiomonas* | 100.0% | 84.3% | EELTRNIPVVMLTARGEEEDRVRGLEVGADDYITKPFSPKELVARIKAVMRRIAPTSVEEVIDVQGLRLDPISHRVTAGE |
| *Providencia* | 100.0% | 81.7% | DSDTRDIPVMMLTARGEEEDRVKGLEVGADDYLIKPFSPKELVARVKAILRRISPMATEDIIEMNGLTLDPTSHRVSSND |
| *Xenorhabdus* | 100.0% | 81.7% | DSSTRDIPVMMVTARGEEEDRVRGLDVGADDYITKPFSPKELVARIKAVLRRVSPMSAEDIINMDGLILDSSSHRVSSQG |
| *Photorhabdus* | 100.0% | 81.2% | DNNVRDIPVMMLTARGEEEDRVRGLEIGADDYITKPFSPKELVARVKAVLRRISPMMIENLIEMDGLILDITSHRVTSQD |
| *Morganella* | 100.0% | 81.2% | DSQLREIPVMMLTARGEEEDRVKGLETGADDYLTKPFSPKELVARVKAILRRISPMAADDLIAMNGLTLDPASHRVTSNE |
| *Vibrio* | 100.0% | 80.3% | EEMTRNIPVVMLTARGEEEDKVRGLEVGADDYITKPFSPKELVARLKAVIRRVTPTALEDVIDVQGLKLDPVSHRVTAND |
| *Proteus* | 100.0% | 79.0% | ENNTRDIPIIMLTAKGEEEDKVKGLETGADDYVIKPFSPKELVARVKAILRRLSPMSAEDVIEFNGLSLDPVSHRVTSQD |
| *Leminorella* | 100.0% | 75.1% | EALTKDIPIIMLTARSEEDDRVQGLESGADDYVTKPFSPKELMARIKAVMRRASPMTEHEQLTFSGLVLDPSSHRVMADD |
| *Arsenophonus* | 100.0% | 74.8% | DQLTRNIPIMMLTARGEEEDRVRGLEIGADDYLTKPFSPKELIARVNAILRRVSAMASDNVIEMNGLKLDPASHRVMSDA |
| *Pseudomonas* | 99.6% | 61.3% | DELTVDIPIIMLTAKGEEDNKIQGLEVGADDYITKPFSPRELVARLKAVLRRTGPGDSEAPIEVGGLLLDPISHRVTIDG |

```
               cov     id
Escherichia    100.0% 100.0%   EPLEMGPTEFKLLHFFMTHPERVYSREQLLNHVWGTNVYVEDRTVDVHIRRLRKALEPGGHDRMVQTVRGTGYRFSTRF
Enterobacter   100.0%  96.5%   NPLDMGPTEFKLLHFFMTHPERVYSREQLLNNVWGTNVYVEDRTVDVHIRRLRKALELSGHDRMVQTVRGTGYRFSTRF
Citrobacter    100.0%  95.6%   NPLDMGPTEFKLLHFFMTHPERVYSREQLLNHVWGTNVYVEDRTVDVHIRRLRKALETSGHDRMVQTVRGTGYRFSTRF
Yokenella      100.0%  95.6%   NPLDMGPTEFKLLHFFMTHPERVYSREQLLNHVWGTNVYVEDRTVDVHIRRLRKALEHSGHDRMVQTVRGTGYRFSARF
Salmonella     100.0%  95.6%   SPLDMGPTEFKLLHFFMTHPERVYSREQLLNHVWGTNVYVEDRTVDVHIRRLRKALEHSGHDRMVQTVRGTGYRFSTRF
Raoultella     100.0%  95.6%   SPLDMGPTEFKLLHFFMTHPERVYSREQLLNHVWGTNVYVEDRTVDVHIRRLRKALEHSGHDRMVQTVRGTGYRFSARF
Klebsiella     100.0%  95.2%   SPLDMGPTEFKLLHFFMTHPERVYSREQLLNHVWGTNVYVEDRTVDVHIRRLRKALEHSGHDRMVQTVRGTGYRFSARF
Cronobacter    100.0%  94.8%   NPLDMGPTEFKLLHFFMTHPERVYSREQLLNNVWGTNVYVEDRTVDVHIRRLRKALEQSGHDRMVQTVRGTGYRFSTRF
Cedecea        100.0%  92.1%   DPLDMGPTEFKLLHFFMTHPERVYSREQLLNNVWGTNVYVEDRTVDVHIRRLRKALETSGHDKMVQTVRGTGYRFSTRF
Hafnia         100.0%  91.7%   APLDMGPTEFKLLHFFMTHPERVYSREQLLNHVWGTNVYVEDRTVDVHIRRLRKALELSGHDKMVQTVRGTGYRFSARF
Serratia       100.0%  91.3%   QPLDMGPTEFKLLHFFMTHPERVYSREQLLNHVWGTNVYVEDRTVDVHIRRLRKALESSGHDKMVQTVRGTGYRFSTRY
Pantoea        100.0%  90.4%   TPLEMGPTEYKLLHFFMTHPERVYSREQLLNHVWGTNVYVEDRTVDVHIRRLRKALEVSGHDRMVQTVRGTGYRFSARY
Brenneria      100.0%  89.5%   LALDMGPTEFKLLHFFMTHPERVYSREQLLNHVWGTNVYVEDRTVDVHIRRLRKALETSGHDKMVQTVRGTGYRFSTRY
Dickeya        100.0%  89.1%   HALDMGPTEFKLLHFFMTHPERVYSREQLLNHVWGTNVYVEDRTVDVHIRRLRKALETSGHDKMVQTVRGTGYRFSTRY
Rahnella       100.0%  89.1%   QALDMGPTEFKLLHFFMTHSERVYSREQLLNNVWGTNVYVEDRTVDVHIRRLRKALETSGHDRMIQTVRGTGYRFSTRY
Pectobacterium 100.0%  89.1%   HALDMGPTEFKLLHFFMTHPERVYSREQLLNHVWGTNVYVEDRTVDVHIRRLRKALETSGHDKMVQTVRGTGYRFSTRY
Erwinia        100.0%  87.8%   QPLDMGPTEYKLLHFFMTHTERVYSREQLLNHIWGTNVYVEDRTVDVHIRRLRKALELSGHDRMVQTVRGTGYRFSARY
Yersinia       100.0%  86.2%   QALDMGPTEFKLLHFFMTHPERVYSREQLLNYVWGTNVYVEDRTVDVHIRRLRKALETDGHDKMVQTVRGTGYRFSTRY
Edwardsiella   100.0%  85.2%   DAVEMGPTEFKLLHFFMTHPERVYSREQLLNHVWGTNVYVEDRTVDVHIRRLRKALEGSGHDRMVQTVRGTGYRFSARF
Plesiomonas    100.0%  84.3%   QPLDMGPTEFKMLHFFMTHQERVYSREQLLNNVWGTNVYVEDRTVDVHIRRLRKALEGSGHDRLIQTVRGSGYRFSTRY
Providencia    100.0%  81.7%   TPIDMGPTEYKLLHFFMTHPERVYSREQLLNYVWGANVYVEDRTVDVHIRRLRKALEAEGHDKMVQTVRGTGYRFSVRY
Xenorhabdus    100.0%  81.7%   QNVEMGPTEFKLLHFFMTHPERVYSREQLLNYVWGTNVYVEDRTVDVHIRRLRKALEVGGHDRMVQTVRGTGYRFSTRY
Photorhabdus   100.0%  81.2%   QVLDMGPTEFKLLHFFMTHPERVYSREQLLNYVWGANVYVEDRTVDVHIRRLRKVLEVGGHDKMVQTVRGTGYRFSTRY
Morganella     100.0%  81.2%   NPLEMGPTEFKLLHFFMTHPERVYSREQLLNYVWGENVYVEDRTVDVHIRRLRKALEADGHDKMIQTVRGTGYRFSVRY
Vibrio         100.0%  80.3%   QPLDMGPTEFKMLHFFMTHQERVYSREQLLNNVWGTNVYVEDRTVDVHIRRLRKALEDAGHDKLIQTVRGAGYRFSTKA
Proteus        100.0%  79.0%   KPIDMGPTEFKLLHFFMTHPERVYSREQLLNYVWGTNVYVEDRTVDVHIRRLRKAIEQDGHDRMIQTVRGTGYRFSARF
Leminorella    100.0%  75.1%   VPLDMGPTEYKLLHFFMSHPERVYSREQLLNQVWGTNVYVEDRTVDVFIRRLRKTLEGSGYDKMLQTVRGSGYRFSTRF
Arsenophonus   100.0%  74.8%   TSIEMGPTEYKLLYFFMTHPERVYNREQLLNYVWGVNVYIEDRTVDVHIRRLRKALAVGGHEKMIQTVRGTGYRFSVRY
Pseudomonas     99.6%  61.3%   KPAEMGPTEYRLLQFFMTHQERAYTRGQLLDQVWGGNVYVEERTVDVHIRRLRKALG-EVYENLVQTVRGTGYRFSTKS
                                     **                 *       *      * **  ** ** *             * **  *
```

**Figure S1. Alignment of PhoB protein sequences used for phylogenetic analysis of binding sites.** PhoB protein sequences from 29 γ-proteobacterial species. Amino acids are color-coded by chemical property; amino acids are only colored if they are identical to the equivalent position in *E. coli*. Asterisks under the alignment indicate amino acids reported to contact DNA (32). The percent coverage (cov) and identity (id) relative to the *E. coli* sequence is indicated next to the genus name.
